# Large-scale phage-based screening reveals extensive pan-viral mimicry of host short linear motifs

**DOI:** 10.1101/2022.06.19.496705

**Authors:** Filip Mihalic, Leandro Simonetti, Girolamo Giudice, Marie Rubin Sander, Richard Lindqvist, Marie Berit Akprioro Peters, Caroline Benz, Eszter Kassa, Dilip Badgujar, Raviteja Inturi, Muhammad Ali, Izabella Krystkowiak, Ahmed Sayadi, Eva Andersson, Hanna Aronsson, Ola Söderberg, Doreen Dobritzsch, Evangelia Petsalaki, Anna K Överby, Per Jemth, Norman E. Davey, Ylva Ivarsson

## Abstract

Viruses mimic host short linear motifs (SLiMs) to hijack and deregulate cellular functions. Studies of motif-mediated interactions therefore provide insight into virus-host dependencies, and reveal targets for therapeutic intervention. Here, we describe the pan-viral discovery of 1,712 SLiM-based virus-host interactions using a phage peptidome tiling the intrinsically disordered protein regions of 229 RNA viruses. We find mimicry of host SLiMs to be a ubiquitous viral strategy, reveal novel host proteins hijacked by viruses, and identify cellular pathways frequently deregulated by viral motif mimicry. Using structural and biophysical analyses, we show that viral mimicry-based interactions have similar binding strength and bound conformations as endogenous interactions. Finally, we establish polyadenylate-binding protein 1 as a potential target for broad-spectrum antiviral agent development. Our platform enables rapid discovery of mechanisms of viral interference and the identification of potential therapeutic targets which can aid in combating future epidemics and pandemics.

## INTRODUCTION

Viruses are obligate intracellular parasites that depend on the host cell machinery for successful infection and replication (Forterre and Prangishvili, 2009). Viruses hijack and deregulate the host cell machinery through virus-host protein-protein interactions (PPIs) that often involve interactions between folded host proteins and viral short linear motifs (SLiMs) (Davey et al., 2011). SLiMs are compact and degenerate protein interaction modules, typically encoded in protein regions between three to ten amino acids in length and often, but not always, found in intrinsically disordered regions (IDRs) of proteins (Elkhaligy et al., 2021; Kumar et al., 2022). SLiM-based hijacking has been reported for all stages of viral infection, including viral cell entry, replication, assembly, release, and subversion of the cellular defense response (Davey et al., 2011; Kadaveru et al., 2008). Mimicry of host SLiMs provides viruses with an elegant solution to the spatial constraints of their genomes as compact SLiM interfaces allow for high functional density within a limited protein region.

Virus-host PPIs have been mapped for several viruses through affinity purification-mass spectrometry (AP-MS) and yeast two-hybrid (Y2H) based approaches (Batra et al., 2018; Davis et al., 2015; Gordon et al., 2020a; Jager et al., 2011; Shah et al., 2018; Shapira et al., 2009). Additionally, more than 200,000 virus-host PPIs have been suggested from computational structure-based pan-viral analyses (Lasso et al., 2019). However, SLiM-based interactions are likely underrepresented in the available large-scale virus-host PPI datasets because the methods used are not optimized to capture low-affinity transient SLiM-based interactions (Benz et al., 2022; Cluet et al., 2020). Consequently, most SLiM-based virus-host PPIs have been identified using low-throughput methods (Kumar et al., 2022). Nevertheless, bioinformatic analysis has suggested that viral mimicry of host SLiMs is a common strategy for viral takeover (Hagai et al., 2014), and many questions remain to be answered by systematic and unbiased pan-viral studies. For example, it is not clear how pervasive the viral use of SLiM-based interactions is, what similarities and differences exist among viral families in terms of preferred host targets, and to what extent virus-host PPIs converge upon specific vulnerabilities in the hosts networks.

In this study, we present an extensive pan-viral dataset of interactions between viral motifs and human protein domains generated by proteomic peptide phage display (ProP-PD) using a phage library containing peptides from 229 RNA viruses and 139 human bait protein domains (Kruse et al., 2021). Based on our results we (i) show that most viruses mimic host SLiMs to interact with host proteins, (ii) identify weak points in cellular pathways that are susceptible to viral interference, (iii) demonstrate that the IDRs of many viral proteins contain multiple overlapping or adjacent SLiMs highlighting high functional density, (iv) show how viral SLiMs can exploit endogenous PPIs by binding host domains with comparable affinities to endogenous ligands, and (v) demonstrate how our approach can identify potential targets for the development of novel antiviral agents.

## RESULTS

### Large-scale screening using an RNA virus peptidome reveals ubiquitous pan-viral SLiM-based mimicry

We screened for virus-host interactions using a phage display library that displays the IDRs from 229 RNA viruses on the major coat protein P8 of the filamentous M13 phage (Kruse et al., 2021). This **Ribo**viria **V**iral **D**isorderome (RiboVD) library (**Table S1**; 19,549 unique 16 amino acid-long peptides; 96.4% confirmed by next-generation sequencing (NGS; **Figure S1**)) contains an almost equal contribution of peptides from positive-sense single stranded RNA ((+) ssRNA) and negative-sense ss RNA ((-) ssRNA) viruses. A minor fraction of the peptides originated from double stranded (ds) RNA viruses and a very small percentage of the peptides from the Hepatitis delta virus which is a circular ssRNA virus (**Figure 1D**). The *Paramyxoviridae* family ((-) ssRNA) contributed most peptides to the library design, followed by the *Coronaviridae* ((+) ssRNA) and the *Rhabdoviridae* ((-) ssRNA) families. Viral families with lesser contribution of peptides were for example *Flaviviridae* ((+) ssRNA; 288 peptides) and *Bornaviridae* ((-) ssRNA; 86 peptides). The differences in the peptide distribution arise from variation in the availability of sequence information for different viral families, as well as length and intrinsic disorder content of the viral proteomes.

**Figure. 1.**
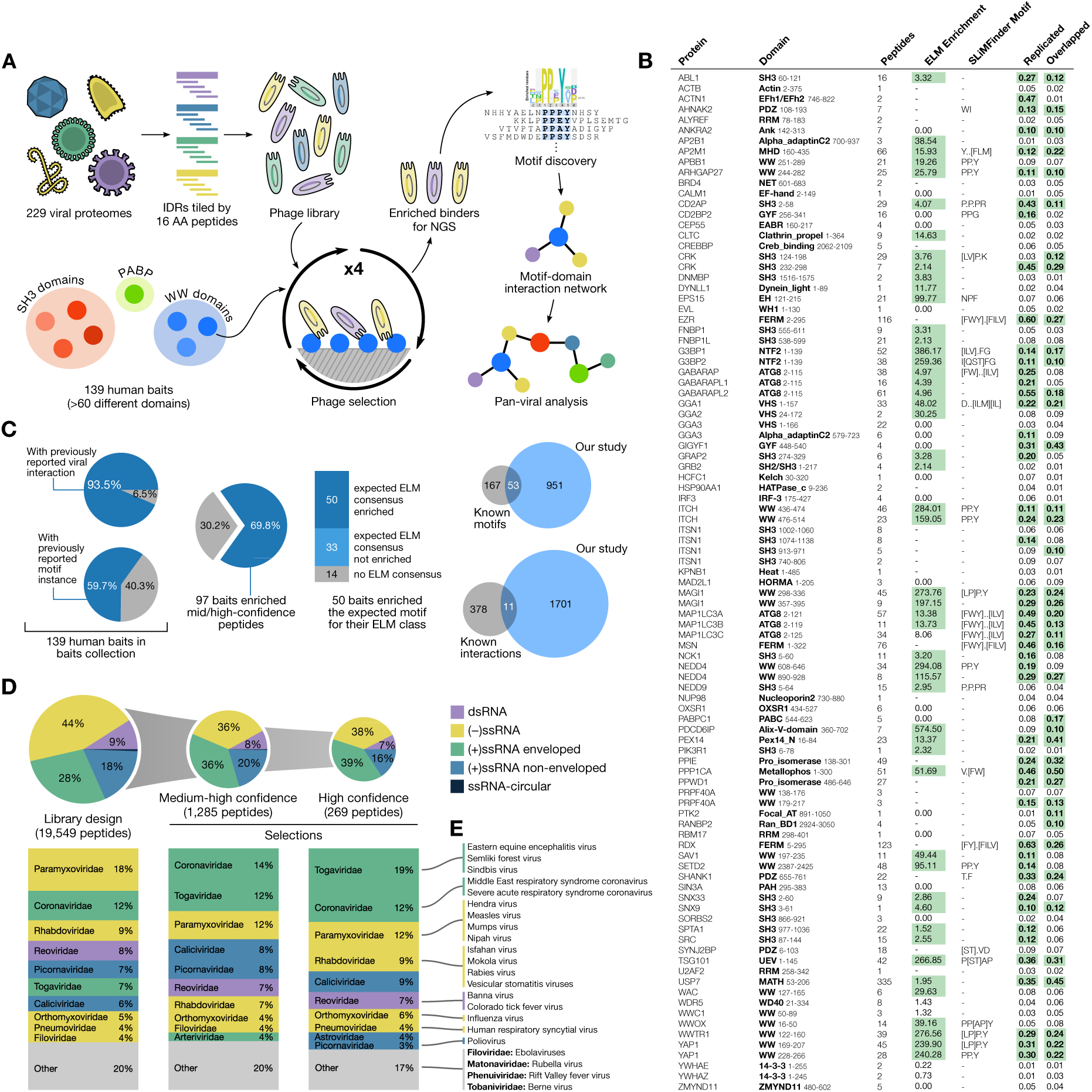
Overview of the RiboVD library design and selection outcome. **(A)** General workflow of the selection and data analysis. **(B)** Overview of the RiboVD selection results showing for each bait the number of enriched medium/high-quality peptides, the enrichment of peptides with sequences matching the consensus motif reported in ELM database (green highlighting indicates a strong enrichment), the de novo generated motifs based on the enriched peptides, and the quality of the selection results (proportion replicated and overlapping peptide – green highlighting indicates a high proportion of peptide). **(C)** Overview of previously the selection results in context of previously known information. Percentage of baits with previously known viral interactors (left, 130, Table S2) and previously reported motif instances (83, Table S4); (center) bait domains that enriched peptides in selections and how their enriched motifs (if any) relate to the 83 previously motif instances; (right) overlap between the RiboVD results and previously reported human-virus SLiM-based interactions (Table S4), or human-virus PPIs (Table S5). **(D)** RiboVD library composition and peptide distribution before and after selections. The representation of peptides for viruses with ssRNA-circular genomes was 103 peptides (0.5%) for the RiboVD library design, and 3 peptides (0.2%) for the medium/high confidence peptide set. **(E)** Examples of representative viruses from different virus families investigated in this study.

Using the RiboVD library, we performed triplicate ProP-PD selections against 139 human bait protein domains (**Figure 1A-B**; **Table S2**), representing more than 60 different domain families. The bait protein domains were mainly selected based on prior reports of interactions with viral proteins (e.g. WW domains (Galinier et al., 2002; Harty et al., 2000), SRC homology 3 (SH3) domains (Arold et al., 1997; Korkaya et al., 2001), and protein phosphatase 1 (PPP1CA) (He et al., 1998)) (**Table S2**). Additionally, we included protein domains that are known to interact with SLiMs (Benz et al., 2022; Teyra et al., 2020) and are involved in cellular processes relevant for viral replication. In total, we identified 1,285 viral peptides binding to 97 domains (**Table S3; Figure 1B**). Notably, these medium/high confidence ligands fulfilled previously defined quality metrics, such as being re-discovered in replicate selections, being highly enriched during selections, and/or containing a consensus motif (Benz et al., 2022). Virus-derived peptides binding to host protein domains were found for nearly 90% of the viral species present in the library, covering all 26 represented viral families. After the selections, there was a shift in the distribution of peptides towards peptides from (+) ssRNA viruses (**Figure 1D**), which may indicate a difference in motif-density between (-) and (+) ssRNA viruses.

To assess the extent to which the RiboVD selections re-discovered known cases of viral motif mimicry we generated a *RiboVD motif benchmarking set* (**Table S4**) which included interactions collected from the Eukaryotic Linear Motif (ELM) database (Kumar et al., 2022), interaction pairs extracted from the Protein Data Bank (PDB), manually curated information from the literature and putative interactions generated by incorporating data from homologous domains. Of 220 viral SLiMs from the benchmarking set that were present in the RiboVD library, 53 were re-discovered by the selections (**Figure 1C**; **Table S4**). The motif -rediscovery rate (24% recall) was high, surpassing our recent benchmarking results against a human disorderome phage library (19.3% rediscovery) (Benz et al., 2022). We further compiled a *virus-host PPI reference set* based on data available in IntAct (Orchard et al., 2014), BioGrid (Oughtred et al., 2021), VirHostNet (Guirimand et al., 2015) and other published sources (**Table S5**). Out of 389 potentially findable virus-host PPIs, only 11 interactions (2.8%) were found by the RiboVD selections. The *virus-host PPI reference set* thus appear largely devoid of SLiM-based interactions.

### Viral motifs bind to common and distinct host targets

The results of the RiboVD selections provided extensive pan-viral information on virus-host PPIs, which allowed us to analyze the relationship between the viral phylogeny and the type of host proteins they interact with (**Figure 2A**). We observed that while some proteins were targeted by specific groups of viral species (*e.g.*, ALYREF RRM and PRPF40A WW by (+) ssRNA viruses), overall the data pointed towards a broad distribution of viral families binding specific baits (*e.g.*, USP7 MATH and WDR5 WD40), indicating large overlaps of the viral SLiM-mediated interactomes.

**Figure 2.**
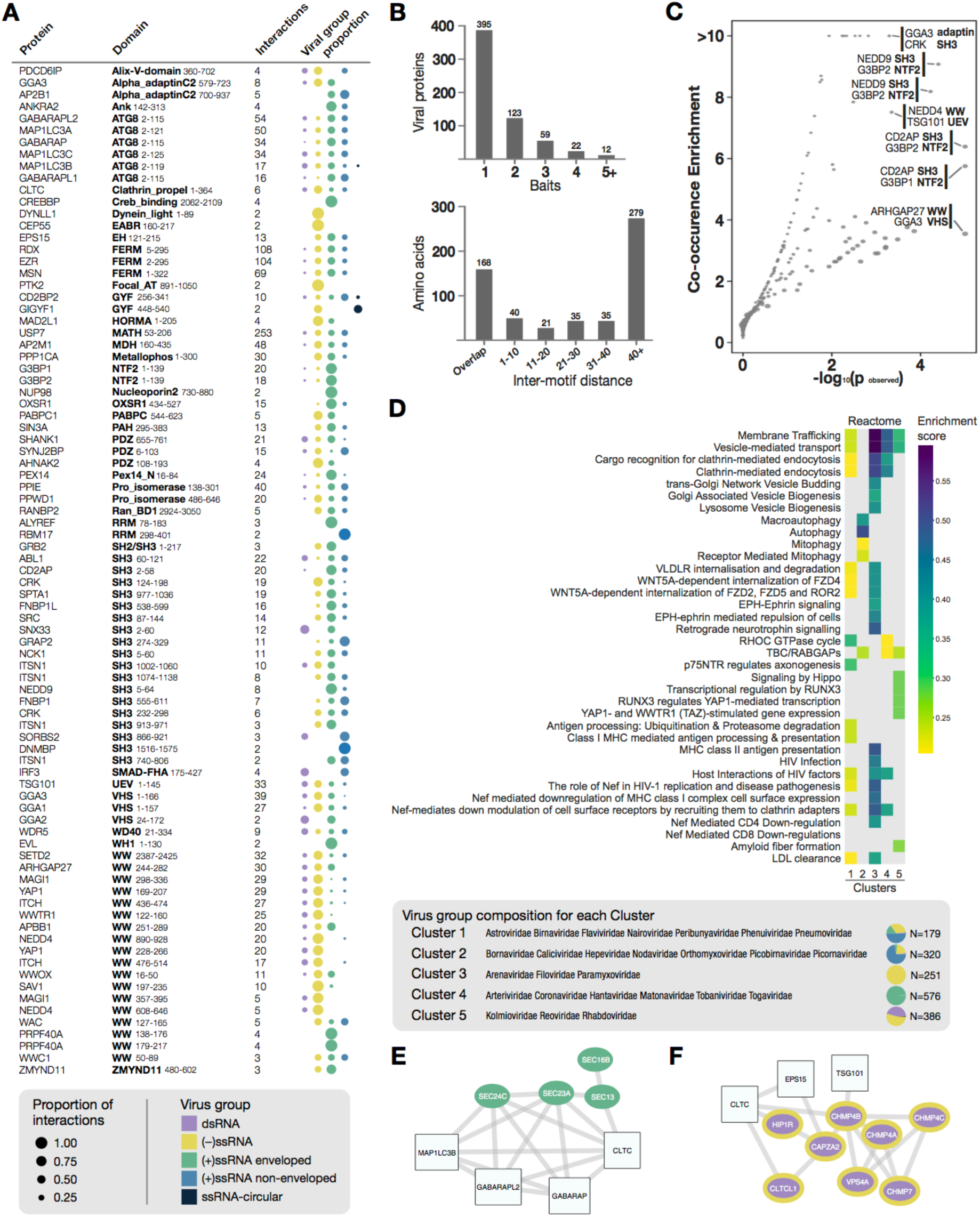
Viral-host PPI and network analysis. **(A)** Overview of the interactions identified per bait, together with the distribution of ligands from different types of viruses. **(B)** Number of screened baits recognized per viral protein (top) and the distance between identified peptides for viral proteins that contain more than one host binding peptide (bottom). **(C)** Analysis of SLiM co-occurrence in viral proteins. **(D)** Clustering of host hijacking network signatures revealed five groups enriched in similar and distinct Reactome pathways. The relative frequency represents the enrichment score adjusted to account for the number of members in each viral family that are contributing to the enrichment. The N indicates the number of identified interactions. **(E)** Sub-network of the COPII complex components (green) identified for cluster 4-based network diffusion, together with their first neighbor bait proteins used in the RiboVD screen. **(F)** Sub-network of the ESCRTIII components (purple with yellow border) identified for cluster 5-based network diffusion, together with their first neighbor bait proteins used in the RiboVD screen.

The results also allowed the exploration of the molecular interplay between distinct types of viral SLiMs (**Figure 2B-C)**. While close to 400 viral proteins bound to a single bait protein, over 200 viral proteins contained more than one type of SLiM. Most motifs found in the same viral protein were distal in the amino acid sequence, suggesting that the interactions with their binding proteins may occur simultaneously. However, 208 out of 578 co-occurring motifs overlapped or were in close proximity (1-10 amino acids), implying that the motifs compete with each other for binding to distinct host proteins **(Figure 2B)**. A subset of SLiMs co-occurred more frequently than would be expected by chance (**Figure 2C**). For example, the NTF2 domains of the Ras GTPase-activating protein-binding proteins 1 and 2 (G3BP1/2) and the SH3 domains of the CD2-associated protein (CD2AP) both interact with co-ocurring SLiMs in the non-structural protein 3 (Nsp3) of several alphaviruses (*Togaviridae)*. Both G3BP1/2 and CD2AP have previously been shown to co-localize with viral replication complexes in alphaviruses (Mutso et al., 2018; Schulte et al., 2016). Moreover, the E3 ubiquitin-protein ligase NEDD4 WW domain (NEDD4 WW) and the tumor susceptibility gene 101 protein UEV domain (TSG101 UEV) binding motifs, which predominantly co-occur in enveloped (-) ssRNA viruses such as Rabies virus (RABV; *Rhabdoviridae*) and Ebola virus (EBOV; *Filoviridae*), enable viral egress by hijacking the endosomal sorting complexes required for transport (ESCRT) machinery (Votteler and Sundquist, 2013). Notably, NEDD4 WW and TSG101 UEV binding motifs frequently co-occurred in close proximity or overlapped, indicating competitive binding, as previously found for EBOV viral matrix protein VP40 (Licata et al., 2003).

### Clustering of host target networks reveals network signatures of viral hijacking

To pinpoint host processes that are commonly targeted by viruses beyond the interactions identified by the RiboVD selections, we used a network diffusion approach. Such analysis assumes that if a human protein is targeted by viral proteins, its neighboring proteins in a protein interaction network are also likely to be important for and/or affected by viral hijacking. Thus, if multiple host proteins fall in a similar region of the network, network modules or signatures relevant to viral hijacking will be highlighted. This analysis allowed us to extract network signature perturbations for each virus in the dataset. Functional enrichment analysis of these signatures revealed that RNA viruses preferentially target proteins involved in protein transport, in particular endocytosis, autophagy, cell morphogenesis, and cell signaling (**Figure S2**; **Table S6**). Next, we searched for network modules or processes that were unique to specific virus types. We clustered the viral families according to their interaction networks and identified five main clusters (**Figure 2D; Figure S3**). While cluster 1 was heterogeneous, the other four clusters were dominated by distinct types of viruses: cluster 2: mostly non-enveloped (+) ssRNA viruses, cluster 3: enveloped (-) ssRNA viruses, cluster 4: enveloped (+) ssRNA viruses, and cluster 5: dsRNA viruses and (-) ssRNA viruses. All viruses except those in cluster 2 targeted processes related to vesicle-mediated transport, with the enveloped (-) ssRNA and (+) ssRNA viruses in cluster 3 and 4 targeting clathrin-mediated endocytosis (**Table S6**; **Figure 2D**). For (-) ssRNA viruses we also observed an enrichment of proteins involved in Golgi associated vesicular budding. In contrast, for the non-enveloped viruses in cluster 2 there was an enrichment of processes associated with autophagy, directly targeting ATG8-like host proteins (microtubule-associated proteins 1A/1B light chain 3 (MAP1LC3s) and gamma-aminobutyric acid receptor-associated proteins (GABARAPs)). The distinct signature for cluster 2 may be related to the fact that non-enveloped viruses do not require trafficking machinery for lytic release but instead use autophagy for non-lytic egress (Bird and Kirkegaard, 2015; Owusu et al., 2021; Sun et al., 2019). Some viruses in cluster 2 such as poliovirus (*Picornaviridae*) have also been reported to use the autophagy machinery during early replication events (Abernathy et al., 2019; Bird et al., 2014).

Overall, there are both similarities and differences in functional enrichments between the different clusters (**Figure 2D**), consistent with hijacking of similar processes but also with distinct signatures of host network interference between different viral groups. For example, comparing the enriched proteins involved in vesicle-mediated transport between the (+) ssRNA viruses in cluster 4 (e.g. *Coronaviridae*) and the (-) ssRNA and dsRNA viruses in cluster 5 (e.g. *Rhabdoviridae* and *Reoviridae*), we found that the former are enriched in proteins linked to the cytoplasmic coat protein complex II (COPII), which sorts cargo from the endoplasmic reticulum (ER) to the trans-Golgi network (Lord et al., 2013), while the latter are enriched in proteins associated with the ESCRT-III complex involved in reverse topology vesicular egress and viral budding (**Figure 2C**; **Figure S4**) (Votteler and Sundquist, 2013). Coronaviruses (in cluster 4) assemble by budding into the lumen of the intermediate compartment at the ER-Golgi interface (Saraste and Prydz, 2021). In contrast, members of the *Rhabdoviridae* family (cluster 5; e.g. RABV and vesicular stomatitis virus (VSV)) bud at the plasma membrane via the ESCRT complex (Votteler and Sundquist, 2013). The result may thus be linked to differences in budding between the distinct viral clusters.

To demonstrate how our RiboVD data can provide deeper insights, we selected protein interactions involved in three biological processes (the ESCRT machinery, endocytosis and protein translation) for detailed investigation.

### Hijacking of the ESCRT machinery highlights motif co-occurrence

Many viruses exploit the ESCRT pathway machinery for viral budding by binding to the TSG101 UEV domain, the WW domains of NEDD4 and the V domain of programmed cell death 6-interacting protein, commonly called ALIX (**Figure 3A**). These interactions facilitate nuclear envelope budding, formation of double-membrane replication complexes and egress of viral particles from the host cell membranes (Vietri et al., 2020; Votteler and Sundquist, 2013). Selections against the three aforementioned ESCRT-related proteins resulted in 81 peptide hits from 12 virus families, most of them from (-) ssRNA viruses. In addition, we identified interactions between the ESCRT associated centrosomal protein of 55 kDa EABR domain (CEP55 EABR) and the Reston ebolavirus (REBOV) nucleoprotein (NP), as well as the RABV protein P (**Figure 3B, G**). CEP55 interacts with TSG101 and ALIX to form a complex that is involved in abscission of the plasma membrane at the midbody during cell division (Lee et al., 2008).

**Figure 3.**
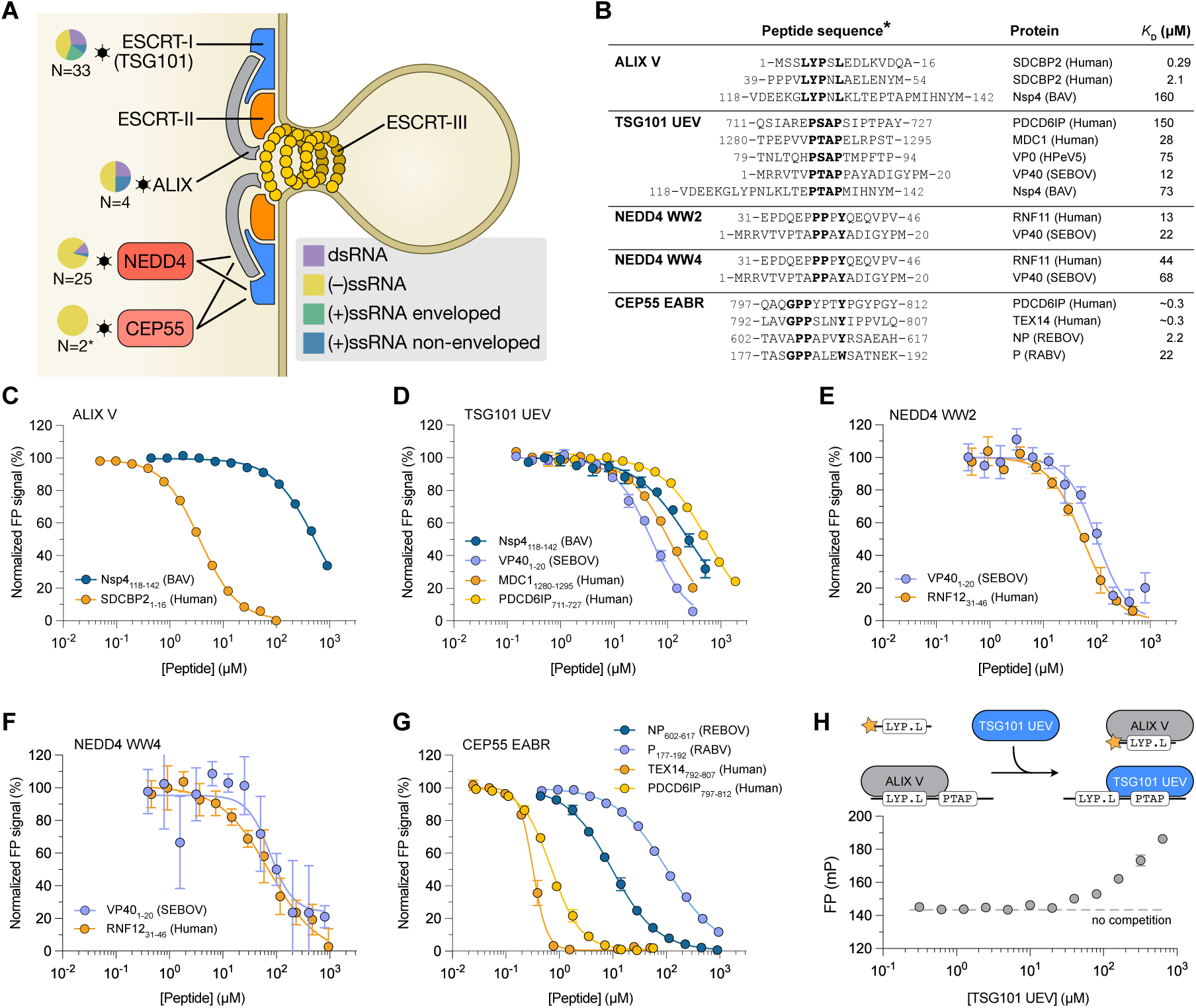
The ESCRT machinery is hijacked by viral SLiMs that bind to NEDD4 WW, TSG101 UEV and ALIX V, and potentially also to CEP55 EABR. **(A)** Schematic representation of the ESCRT pathway leading to reverse topology budding. Pie charts next to each target show the class of viral species hijacking it. N represents the number of identified interactions. **(B)** Overview of the peptides that bind to ESCRT pathway proteins for which the affinities were measured. Residues constituting the recognition motif are shown in bold. (**C-G**) FP-monitored displacement experiments of viral and human peptides, and ESCRT proteins. Data are represented as normalized means ± SD. For detailed information on the peptides used in this study see Table S7. **(H)** FP-monitored displacement experiment of Nsp4_118-142_ (BAV) shows that the interaction with ALIX V and TSG101 UEV is mutually exclusive.

We determined affinities for ALIX V, NEDD4 WW, TSG101 UEV and CEP55 EABR with viral and human peptides using a fluorescence polarization (FP) based assay (**Figure 3B-G**; **Figure S5; Table S7**). The affinities of the viral SLiMs for their respective protein domains were found to be in the low-to-mid micromolar range (**Figure 3B**), which is typical for SLiM-based interactions (Benz et al., 2022; Ivarsson and Jemth, 2019). Viral and endogenous host SLiMs bound with comparable affinities to NEDD4 WW and TSG101 UEV domains. In contrast, the viral ALIX V domain ligand Nsp4_118-142_ (BAV) showed a >300-fold weaker affinity compared to the endogenous ligand derived from syntenin-2 (SDCBP2_1-16_ (Human)) (**Figure 3B**). Similarly, the viral CEP55 EABR peptide binders were found to bind the protein with one to two orders of magnitude weaker affinity than the endogenous ligands **(Figure 3G)**. A higher concentration of the viral ligands would hence be necessary to outcompete the endogenous interactions.

Following up on co-occurring motifs, we noted a close proximity of the ALIX V binding LYPNL motif and the TSG101 UEV binding PTAP motif in Nsp4 of Banna virus (Nsp4_118-142_ (BAV)) (**Figure 3B**). We therefore investigated whether the four amino acids separating the two motifs were sufficient to allow simultaneous interaction of both domains with Nsp4_118-142_ (BAV) or if there is competition between the two binding motifs. We challenged a pre-formed complex of ALIX V domain and Nsp4_118-142_ (BAV) peptide with increasing concentrations of TSG101 UEV in the presence of a constant concentration of fluorescein isothiocyanate (FITC)- labeled ALIX V-binding peptide (FITC-gag_493-502_ (HIV1); **Figure 3H**). The observed increase in FP signal with increasing concentrations of TSG101 UEV supported a model of mutually exclusive binding of the TSG101 UEV and ALIX V domains, providing a validated example of competitive binding between the two distinct adjacent motifs. Intriguingly, the Banna virus lacks a membrane envelope but could use the ESCRT pathway for non-lytic viral egress or for the formation of double-membrane replication factories as described for the related bluetongue virus (Wirblich et al., 2006).

Overall, our results for the ESCRT pathway support and complement previous findings and suggest that competitive binding of motifs in close proximity allows viruses to temporally regulate host protein hijacking.

### RiboVD screening reveals hijacking of clathrin adaptors

Viruses frequently mimic SLiMs that bind to proteins involved in the endocytic trafficking machinery (**Figure 2**; **Figure S3**). These interactions involve clathrin (discussed in the following section) or its adaptors (**Figure 4A**). We validated interactions with the Mu homology domain (MHD) of the AP-2 subunits mu (AP2M1 MHD), which is involved in cargo selection and endocytic vesicle formation at the plasma membrane, and with the GAE and the VHS domains of the ADP-ribosylation factor-binding protein GGA3 (GGA3 GAE and GGA3 VHS), involved in cargo recognition and trafficking between the trans-Golgi network and endosomes (Bonifacino, 2004; Puertollano et al., 2001) (**Figure 4B-E**). We found that the interactions of the AP2M1 MHD, GGA3 GAE and GGA3 VHS domains (**Figure 4B-D**) with viral motifs were in the low-to-mid micromolar range, and that they bound with similar, or lower affinities than the tested endogenous interactions similar to the ESCRT pathway interactions described above (**Figure 4B-E, G; Figure S5; Table S7**). We further validated that the nucleoprotein (NP) from Zaire ebolavirus (ZEBOV) has both a _340-_**Y**QQ**L**_-343_ motif and a _466-_**Y**GE**Y**_-469_ motif that that bind to AP2M1 MHD and GGA3 GAE domain, respectively, with low micromolar affinity (**Figure 4C, E**). The interactions between GGA3 VHS and AP2M1 MHD and full-length NP (ZEBOV) were confirmed by glutathione transferase (GST)-pulldown experiments (**Figure 4F; Table S8**). Finally, we confirmed that the interaction between NP (ZEBOV) and GGA3 GAE is motif dependent, as the interaction was lost upon motif mutation (NP ZEBOV mut 1: Y469A). In contrast, the AP2M1 interaction was retained despite two mutations in the AP2M1 binding motif (NP ZEBOV mut 2: Y340A/L343A). Inspection of the NP sequence revealed six potential AP2M1 binding motifs (YxxΦ), all of which may contribute to binding (**Figure S6**). These results corroborate previous findings linking the ebolavirus NP to clathrin adaptor hijacking (Batra et al., 2018; Garcia-Dorival et al., 2016), and illustrate how a single viral protein can exploit different parts of endocytic trafficking by mimicking different trafficking motifs.

**Figure 4.**
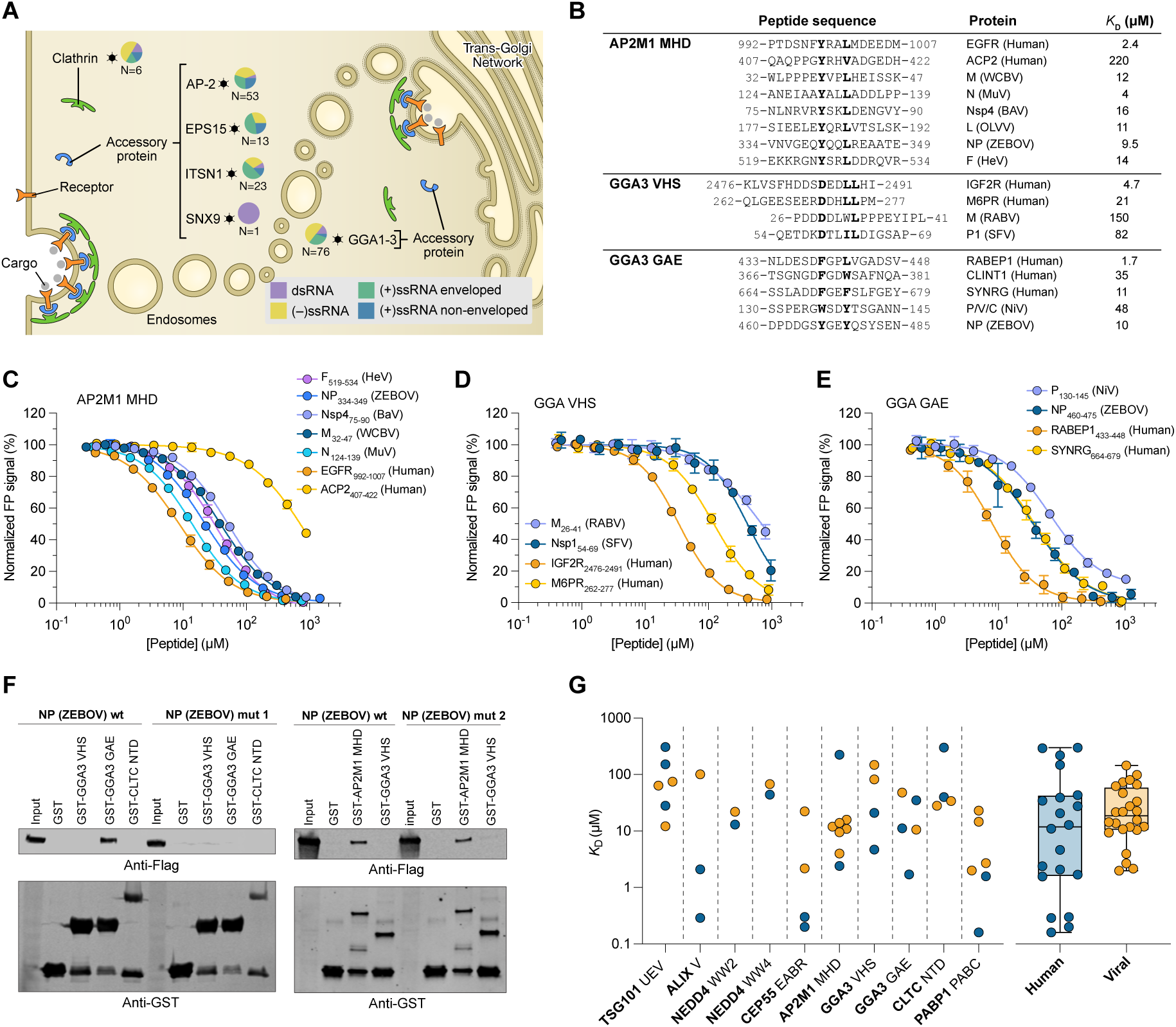
Viral mimicry of distinct trafficking motifs binding to clathrin adaptors. **(A)** Schematic representation of clathrin adaptor vesicle coat components for which viral ligands were found: AP2B1 and AP2M1 (collapsed as AP-2), CLTC, EPS15, ITSN1, SNX9, GGA1, GGA2 and GGA3 (collapsed as GGA1-3). Pie charts show the class of viral species hijacking the domain. N indicates the number of identified interactions. **(B)** Overview of affinity data for peptides interacting with clathrin adaptor proteins. (**C-E**) FP-monitored affinity measurements of viral and human peptides and host proteins. Data are represented as normalized means ± SD (n ≥ 3). **(F)** Capture of full-length viral proteins by GST-tagged domains as visualized by Western blot. The interaction between NP (ZEBOV) and GGA3 GAE is lost upon motif mutation (Y469A). NP (ZEBOV) also interacts with AP2M1 MHD, but the introduced motif mutations (Y340A/L343A) did not abrogate binding suggesting additional AP2M1-binding SLiMs in NP (ZEBOV). **(G)** Overview of all affinity data generated for viral (orange) and host (blue) peptides generated in this study

### Eastern equine encephalitis virus Nsp3 interacts with the N-terminal domain of clathrin and blocks receptor trafficking

Next, we focused on viral mimicry of clathrin binding motifs. The N-terminal domain of clathrin (CLTC NTD) is a β-propeller repeat that binds SLiMs through four different binding sites (Muenzner et al., 2017; Willox and Royle, 2012) (**Figure 5G**). Our selection revealed three viral peptides containing the classical clathrin box motif (LΦxΦ[DE]): a previously described motif in the mu-NS protein of Reovirus type 1 (MRV1) (Ivanovic et al., 2011) together with novel motifs in the Nsp3 of the highly pathogenic Eastern equine encephalitis virus (EEEV) and in the RNA-directed RNA polymerase of the Seneca Valley virus. We confirmed the motif-dependent interaction between the Nsp3_1765-1780_ (EEEV) peptide and clathrin by FP affinity measurements and GST pulldown experiments (**Figure 5A-B, E; Figure S5; Table S8**). We further demonstrated, by an *in-situ* proximity ligation assay (PLA), that the interaction between endogenous clathrin and FLAG-tagged full-length Nsp3 (EEEV) can occur in a cellular setting, mediated by the identified motif (_1771_-**L**IT**FD**-_1775_) (**Figure 5C**; **Figure S7**).

**Figure 5.**
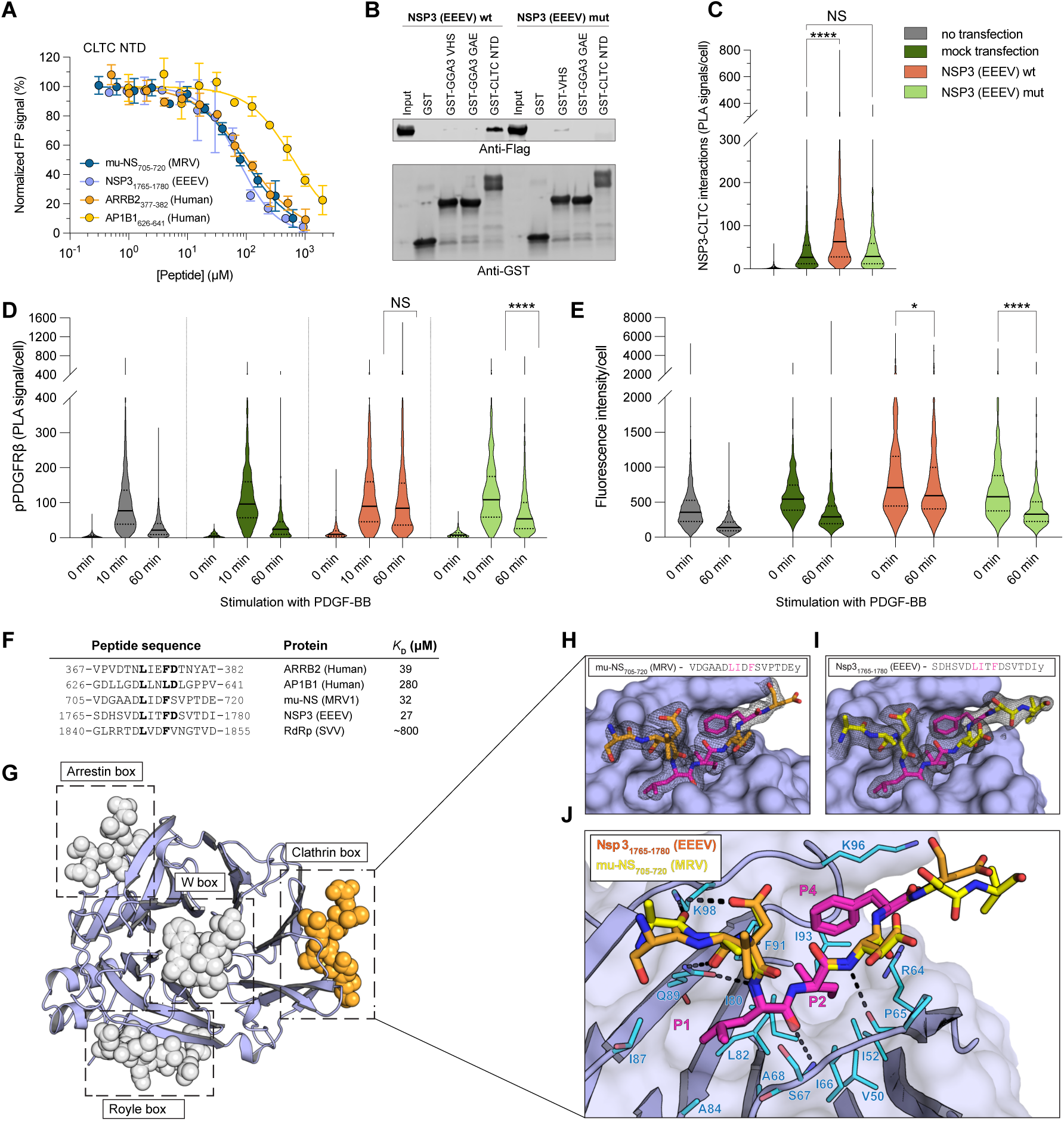
The Nsp3 (EEEV) cathrin box motif is responsible and sufficient for the interaction with clathrin and facilitates the disruption of native cell trafficking. **(A)** FP-monitored affinity measurements of viral and human peptides binding to CLTC NTD. Data are represented as normalized means ± SD (n ≥ 3). **(B)** Capture of full-length Nsp3 (EEEV) by GST-tagged CLTC NTD visualized by Western blot. The interaction is lost upon SLiM mutation (Nsp3 (EEEV) mut; F1774A/D1775A). **(C)** The interaction between clathrin and full-length Nsp3 (EEEV) in HEK293T cells probed by PLA. The results show PLA signal per cell measured in over 1000 cells per construct in six biological replicates and are presented as violin plots with indicated median and interquartile range. P (HeV) mut was used as a mock transfection. Significance was determined by Kruskal-Wallis rank sum test with Dunn’s test and Bonferroni correction as a post-hoc test to compare all groups; \**p*< 0.05, \*\**p*< 0.01, \*\*\**p*<0.001, and \*\*\*\**p*<0.0001. **(D)** Activation of pPDGFRβ (phosphorylation of Y751) in HEK293-PDGFRβ-HA cells probed by PLA. After activation with PDGF-BB, receptor phosphorylation persisted after 60 minutes in the presence of Nsp3 (EEEV) indicating disturbed clathrin-mediated endocytosis. Results were quantified as in **C** for more than 1000 cells per construct over three (mock transfection) or six (Nsp3 (EEEV) wt, Nsp3 (EEEV) mut and no transfection) biological replicates. Significance was determined as in **C**. Corresponding fluorescence microscopy images for **C** and **D** are shown in **Figure S10-11**. **(E)** Quantification of the retention of activated PDGFRβ at the plasma membrane as observed by cell surface fluorescence assay. Integrated fluorescence intensity was measured for more than 300 cells per construct over 3 biological replicates. Significance was determined by Mann-Whitney test with Bonferroni correction for pairwise comparison. **(F)** Alignment of peptides binding to CLTC NTD, with corresponding affinities. **(G)** Structure of CLTC NTD with four motif-binding sites and bound peptides shown as gray spheres (coordinates for peptides bound to Arrestin, Royle and W boxes were obtained from the PDB entries 1UTC and 5M5T). The peptide bound to the clathrin box is colored orange and represents the binding site of the viral peptides investigated in this study. **(H-J)** Crystal structure of short linear motifs from Nsp3 (EEEV) and mu-NS (MRV1) bound to the clathrin box of CLTC NTD. Panels **H** and **I** show the bound peptides (colored sticks) with the corresponding electron density maps calculated using the final model (black mesh). Conserved motif residues are shown in magenta. (**J)** Overlay of the two peptides highlighting the conserved binding mode. The CLTC NTD residues engaged in interaction with the viral protein-derived peptides are shown as blue sticks and hydrogen bonds are highlighted by dotted lines. Coloring of the peptide ligands is the same as in **H** and **I**. The numbering of the peptide residues starts with P1 being the first position of the consensus recognition motif, while the first residue before the motif is numbered P-1.

To further characterize the interactions with clathrin, we solved the structure of CLTC NTD co-crystallized with either Nsp3_1765-1780_ (EEEV) or mu-NS_705-720_ (MRV1) (**Figure 5F-I; Table S9**). In both complexes, the structure of the CLTC-NTD was nearly identical, with a root mean square deviation of less than 0.3 Å, and the central eight residues of the peptides well defined in the electron density (**Figure 5G-I**). The viral peptides bound exclusively to the hydrophobic clathrin box binding pocket, located between blade one and two of the N-terminal β-propeller domain. Structural comparison of the bound viral peptides with an available structure of the host ligand AP2B1 (PDBid: 5M5R**; Figure S8**) (Muenzner et al., 2017) revealed a similar placement of corresponding residues in the hydrophobic pocket.

The structures supported viral mimicry of the clathrin box motif and a direct competition between viral and human clathrin-binding proteins, which suggested potential interference of Nsp3 (EEEV) with the normal function of clathrin. To explore this competition we used the platelet-derived growth factor receptor β (PDGFRβ) as a model for a receptor tyrosine kinase that is internalized via clathrin-mediated endocytosis (Goh and Sorkin, 2013). After activation by its ligand PDGF-BB, the receptor is phosphorylated at several residues in the cytoplasmic part, internalized primarily via clathrin-mediated endocytosis (Rogers and Fantauzzo, 2020), and subsequently degraded (**Figure S9**). We hypothesized that the binding of Nsp3 to clathrin would interfere with clathrin-mediated endocytosis resulting in impaired internalization of activated PDGFRβ. We observed a sharp increase in PLA signal probing for activated PDGFRβ phosphorylated at Tyr751 (Heldin et al., 2019) 10 minutes after activation with PDGF-BB in all four experimental setups. Consistent with our hypothesis, the signal decreased after 60 minutes in non-transfected cells, mock-transfected cells, or in cells transfected with a motif-mutant construct Nsp3 (EEEV) mut but persisted in cells transfected with wild-type Nsp3 (EEEV). (**Figure 5D; Figure S10**). The clathrin-Nsp3 (EEEV) interaction thus interferes with normal receptor signal attenuation. To confirm that the activated receptor remained on the cell surface, we performed a cell surface fluorescence assay, which confirmed the presence of PDGFRβ on the plasma membrane 60 minutes post stimulation, when cells were transfected with Nsp3 (EEEV) wt but not when they were treated with other control constructs **(Figure 5E; Figure S11)**, further supporting the notion that the Nsp3 (EEEV) interferes with normal clathrin-mediated endocytosis. Importantly, we here used PDGFRβ as a model system, but the results suggest a more general inhibition of clathrin-dependent trafficking. The clathrin-Nsp3 (EEEV) interaction could disrupt surface display of receptors in an analogous manner to HIV1 Nef (Kwon et al., 2020), or alternatively serve to recruit clathrin to viral replication centers, as previously shown for the clathrin-mu-NS (MRV1) interaction (Ivanovic et al., 2011). The exact outcomes of viral clathrin hijacking may warrant further exploration.

### The C-terminal domain of the polyadenylate-binding protein 1 is a target of viral hijacking

In order to successfully replicate, viruses need to hijack the host translational machinery (Bushell and Sarnow, 2002). While our screen did not reveal enrichment of interactions with translational machinery proteins, we identified a number of viral peptides that bind to the C terminal domain of polyadenylate-binding protein 1 (PABP1 PABC). PABP1 normally binds to the poly(A) tail of mRNA, stabilizing it and promoting translation initiation (**Figure 6A**) (Mangus et al., 2003; Smith et al., 2014). PABP1 is commonly degraded by viral proteases to repress translation of endogenous proteins, but can also be subjected to viral hijacking to promote translation of viral proteins (Lei et al., 2021; Smith and Gray, 2010). Using the PABC domain of PABP1 as a bait, we uncovered interactions with three viral peptides that contain a typical PABP-interaction motif (**Figure 6C**). The peptides were found in the non-structural protein P/V/C of the highly pathogenic Hendra virus (HeV; P/V/C_183-198_ (HeV)), and in the nucleoprotein (N) of human coronavirus 229E (N_351-366_ (HCoV 299E)) and Berne virus (N_2-17_ (BeV)). The PABP binding motif in HeV is also conserved in the closely related Nipah virus (NiV) (P/V/C_183-198_, **Figure 6C**). We determined the affinities of PABP1 PABC for four peptides from HeV, NiV, HCoV 229E and BeV (**Figure 6B**). The P/V/C_183-198_ (HeV) peptide and the N_2-17_ (BeV) were the highest affinity viral PABC ligands. They bound with similar affinity as the endogenous ligand PABP-interacting protein 1 (PAIP1_125-140_), but ten-fold weaker than the peptide from the endogenous PABP inhibitor PABP-interacting protein 2 (PAIP2_108-123_) **(Figure 6B-C**). The motif-dependent interactions with PABP1 PABC were validated with full-length N (HCoV 299E) and P (HeV) by GST-pulldown (**Figure 6D**).

**Figure 6.**
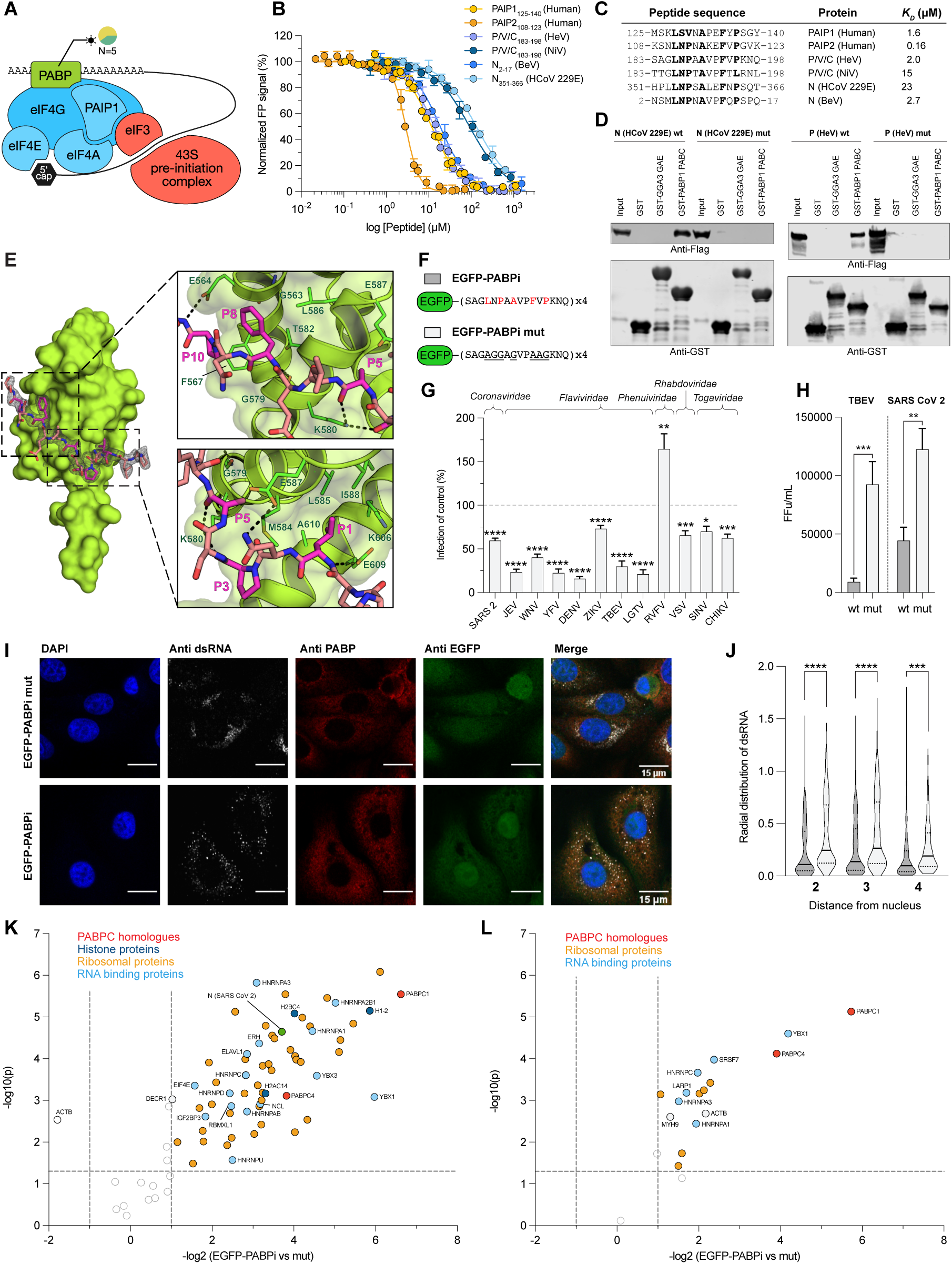
PABP1 is subjected to viral interference and serves as a valid target for broad-spectrum antiviral inhibition. **(A)** Schematic representation of the “closed loop” structure, that promotes translation initiation and ribosomal subunits recruitment. The components of eIF4F are shown in blue and the pre-initiation complex in red. The pie chart next to PABPC1 shows the class of viral species that hijack it. N indicates the number of identified interactions. **(B)** FP-monitored displacement experiments of viral and human peptides and the PABC1 PABC domain. Data are represented as normalized means ± SD (n ≥ 3). **(C)** Alignment of the human and viral peptides interacting with PABP1 PABC. The residues constituting the recognition motif are shown in bold. **(D)** Interactions between full-length viral proteins and GST-tagged PABP1 PABC visualized by western blot. The interaction was lost upon mutation of the PABC interaction motif in viral proteins N (HCoV 229E) mut and P (HeV) mut. **(E)** Structural model of the PABC domain bound to viral peptide N_351-366_ (HCoV 229E) with corresponding electron density map (black mesh). The two side-panels show the close up of two main binding pockets that facilitate interaction with hydrophobic residues in position P1 and P8 of the peptide. The residues responsible for motif recognition are colored purple. **(F)** Schematic representation of EGFP-PABPi and the negative control (EGFP-PABPi mut) lentiviral constructs. **(G)** The antiviral effect of EGFP-PABPi against a selection of different RNA viruses compared with EGFP-PABPi mut. VeroE6 or VeroB4 cells were transduced with EGFP-PABPi or EGFP- PABPi mut lentivirus and subsequently infected with the respective virus. The antiviral effect was measured by percentage of infected cells in EGFP-PABPi compared to EGFP-PABPi mut. **(H)** Corresponding viral titers in the supernatant of cells infected by TBEV and SARS-CoV-2 as determined by focus forming (FFu) assay (**I**) Representative confocal microscopy images of VeroB4 cells transduced with EGFP-PABPi or EGFP-PABPi mut and infected with TBEV (multiplicity of infection (MOI) 1) after 24 hours. (**J**) Quantification of the radial coefficient of variation (RadialCV) of dsRNA intensity in the different fractions of the cell. Each dot represents one infected cell. (**K-L)** Mass spectrometry analysis of differential expression in lentivirus transduced cells expressing EGFP-PABPi or EGFP-PABPi mut and infected with SARS-CoV-2 (VeroE6; **K**) or TBEV (VeroB4; **L**).

To determine the binding mode of the viral peptides, we attempted to co-crystallize the PABP1 PABC domain with the P/V/C_183-19 8_ (HeV) or the N_351-366_ (HCoV 299E) peptides. The PABC-N_351-366_ (HCoV 229E) complex crystallized readily, and the structure was solved to 1.93 Å resolution. In the complex, the peptide is bound in an extended conformation spanning over two hydrophobic PABC pockets located between helices α2 and α3, as well as α3 and α5, respectively (**Figure 6E**). Alignment of the PABP1 PABC binding peptides showed recurrence of a Leu residue at position 1 and of a hydrophobic residue at position 8 (**Figure 6C**), which is a Phe in the N_351-366_ (HCoV 229E) peptide. The structure of PABC-N_351-366_ (HCoV 229E) revealed that the conserved Leu at P1 and Phe at P8 sit in deep hydrophobic pockets which were previously describe to be essential for binding of PAIP2 (Kozlov and Gehring, 2010) and PAIP1 (Munoz-Escobar et al., 2015). A comparison of the binding of N_351-366_ (HCoV 229E) and the human PAIP1 peptide (PDBid 3NTW) revealed a very similar molecular arrangement with a root mean square deviation of < 0.4 Å (**Figure S8**). These results support direct competition between the viral and endogenous PABC ligands.

### The PABC-binding HeV peptide acts as a broad-spectrum inhibitor of viral replication

We reasoned that targeting PABP1 using a PABC-binding peptide could be used to inhibit viral replication of those viruses that rely on PABP1 for efficient translation. For example, the Nsp3 protein from Severe acute respiratory syndrome coronavirus 2 (SARS-CoV-2) interacts with the PABP1 ligand PAIP1 to form a ternary complex with PAIP1 and PABP1, which stimulates viral protein translation (Lei et al., 2021). We generated a lentiviral construct expressing four copies of the P/V/C_183-198_ (HeV) peptide N-terminally fused to EGFP (EGFP-PABPi) and tested its ability to inhibit infection of a panel of RNA viruses (**Figure 6G**). EGFP-PABPi reduced the infection level of almost all viruses tested, with the exception of the Rift Valley fever virus (RVFV). The stimulatory effect on RVFV infection by EGFP-PABPi may be related to a previous finding describing the necessity for the RVFV to sequester PABP1 in nuclear speckles for efficient replication (Copeland et al., 2013). An inhibitory effect of EGFP-PABPi was further demonstrated by low viral titers of the Tick-borne encephalitis virus (TBEV) and SARS-CoV-2 as compared to the control (**Figure 6H)**. To analyze how the presence of the EGFP-PABPi affected the viral replication complex in TBEV infected cells, we detected the viral dsRNA produced within these complexes. We found that the presence of EGFP-PABPi resulted in a more diffuse distribution of the viral replication complexes (**Figure 6I**). The lower concentration and altered localization of replication complexes could explain the lower viral infectivity, although the exact details of how EGFP-PABPi perturbed the viral infection remain to be elucidated. The results support the notion that targeting the peptide binding pocket of PABC blocks replication of a broad panel of RNA viruses.

Finally, we evaluated the specificity of the EGFP-PABPi peptide for its target in human (HEK293, uninfected) or green monkey cells (TBEV infected VeroB4 or SARS-COV-2 infected VeroE6) by AP-MS experiments. Consistent with our results, EGFP-PABPi pulled down PABPC1, together with its homolog PABPC4 in both uninfected HEK293 and virus-infected VeroB4 or VeroE6 cells (**Figure 6K-L; Figure S13; Table S13**). The PABP1/4 proteins were pulled down together with several RNA-binding proteins and with ribosomal proteins, in line with the association of PABP1/4 with the mRNA processing and translation machinery. From cells infected with SARS-CoV-2, EGFP-PABPi additionally pulled down the viral N protein, and its human ligand G3BP1 (Kruse et al., 2021). Overall, the analysis confirmed that the EGFP- PABPi is specific for PABC domain-containing proteins and can successfully be used to attenuate viral replication in a pan-viral manner.

## DISCUSSION

In this study, we present a large-scale pan-viral assessment of how viruses use SLiM-based mimicry to bind host proteins and outcompete endogenous interactors. In total, we found 1,712 virus-host PPIs involving 679 viral proteins from 233 viral species, and 97 globular domains from 87 human proteins, yielding an unprecedented, multilayered dataset on virus-host PPIs. We found that all RNA virus families included in this study have SLiMs that can interact with host proteins. Our results attempt to fill the gaps in host-pathogen interactomes generated by other experimental approaches (e.g., AP-MS and Y2H), with the added value of providing information about the binding motifs with amino acid resolution.

At the highest level, the results give an overview of processes that are frequently targeted by viruses of different families (**Figure 2**). As expected, we found that endocytic transport is a common target of viral hijacking and that different parts of the endocytic machinery are targeted by different viruses with distinct classes and combinations of SLiMs. Closer examination of the data revealed both common strategies of viral hijacking used by unrelated viruses as well as distinct features even among closely related viruses, as demonstrated by the heterogeneous clustering of viruses.

At the molecular level, the results provide exact interaction interfaces in viral proteins. This detailed information can be used to reveal the concerted action of co-occurring motifs in the targeting of human proteins as well as instances of motif competition. We found that adjacent or overlapping SLiMs are common in the IDRs of viral proteomes and likely compete for binding to their host targets **(Figures 2-3)**. Such mutually exclusive binding could provide temporal control that ensures successful hijacking of vital pathways at the appropriate time in the infection process. Closely located or overlapping WW and TSG101 binding motifs are also found in human proteins such as SIMPLE (Lee et al., 2012; Ludes-Meyers et al., 2004), suggesting that competing motifs interacting with the ESCRT machinery are not unique to viruses but represent a more general regulatory approach.

To gain a deeper understanding of the binding and function of viral SLiMs we analyzed the affinity of 25 virus-host PPIs for ten human protein domains and solved the crystal structures of three complexes. Our results showed that the viral ligands bind to the same binding sites as the host ligands and thereby may inhibit host processes, as shown for clathrin-binding Nsp3 (EEEV) (**Figure 5**). In contrast to some of the previous literature (Davey et al., 2011), the viral ligands did generally not exhibit higher affinities for the human targets (**Figure 4G**). The affinities of both host-host and viral-host PPIs cover a wide range of over more than three orders of magnitude with no clear pattern as to which has the higher affinity. The key to efficient hijacking by the viral ligands described here might be found in the high local concentration of viral proteins that are generated in virus infected cells, which is particularly relevant to interactions occurring late during the viral life cycle (e.g. ESCRT pathway). The PABP-binding HeV peptide is an interesting case, as it binds its target with similar affinity to the host ligand PAIP1, a co-activator of translation, but, both bind tenfold weaker than the engogenous PABP inhibitor PAIP2. Thus the affinities of both viral and host ligands appear to be tuned to the functional role of the interaction (transient binding, or blocking of the target).

Given the omnipresent risk of new emerging viruses, there is an urgent need to systematically map virus-host PPIs. We have shown that the PABP1 PABC domain can be targeted to block viral replication in a pan-viral manner. This demonstrates that the identification and targeting of SLiM-based virus-host PPIs may be a viable strategy for the development of novel antiviral drugs. Previous examples of inhibition of viral infection by targeting human proteins include for example targeting of the interaction between the ebolavirus protein VP30 and host protein PP2A-B56 (Kruse et al., 2018), and inhibition of the interaction between N (SARS-CoV-2) and human G3BP1/2 (Kruse et al., 2021). Exploring host proteins as drug targets instead of their viral counterparts is attractive because it has proven more difficult for the virus to evolve resistance to such antiviral agents (Gordon et al., 2020b; Lin and Gallay, 2013). In addition, the same host proteins or host processes are often targeted by a variety of different viruses, which opens new avenues for the development of pan-viral inhibitors, which will contribute towards our preparedness against emerging viral threats (Bekerman and Einav, 2015).

In conclusion, we show that SLiM-based hijacking of host proteins is widespread among RNA viruses. Our data contribute to a better understanding of the molecular details of host cell subversion, and pinpoint novel targets for innovative inhibitor design. Despite the scale of this analysis, we have only started to tap into the host proteins that are targeted by viruses. In the future, we envision studying an even larger collection of bait proteins. We believe that our study will be valuable to molecular virologists refining the mechanistic understanding of viral infections and that pan-viral data will facilitate the search for novel broad-spectrum inhibitors for use against existing and novel emerging viruses.

## MATERIALS AND METHODS

Reagents and resources are summarized in **Table S11**.

### Recombinant protein expression and purification

Proteins (**Table S2**) were expressed in *E. coli* BL21(DE3) as GST-tagged proteins in 2YT growth media (16 mg/mL peptone, 10 mg/mL yeast extract, 5 mg/mL NaCl) supplemented with appropriate antibiotics (50 µg/mL kanamycin (Kan) for pETM33 constructs and 100 µg/mL ampicillin (Amp) for pHH1003 constructs) at 37°C. After reaching an OD_600_ of 0.6 protein expression was induced with 1 mM isopropyl β-D-1-thiogalactopyranoside (IPTG). Proteins were expressed either for 4 hours at 30°C or overnight at 18°C. Bacterial cultures were harvested by centrifugation (4,500 g, 10 minutes) at 4°C and resuspended in lysis buffer A (PBS supplemented with 1% Triton, 10 µg/mL DNase I, 5 mM MgCl_2_, 10 µg/mL of lysozyme, and cOmplete™ EDTA-free Protease Inhibitor Cocktail (Hoffman-La Roche) when the protein was used for phage display selections, or in lysis buffer B (50 mM Tris/HCl pH 7.8, 300 mM NaCl, 10 µg/mL DNase I and RNase, 4 mM MgCl_2_, 2 mM CaCl_2_ and cOmplete EDTA-free Protease Inhibitor Cocktail) when the protein was used for FP affinity determination experiments. Cells were lysed either with two cycles of 20 seconds sonication with 2 seconds pulses, or with a cell disruptor apparatus at 1.7 kBar. The lysate was clarified by centrifugation (20,000 g, 40 minutes) and the supernatant was filtered through a 0.2 µm sterile PES filter, transferred to Pierce Glutathione Agarose and purified according to the manufacturer’s protocol. For proteins used in FP experiments additional purification steps were performed. After elution, the His/GST tag was enzymatically cleaved with either Thrombin or PreScission protease overnight at 4°C. The sample was then applied to a nickel Sepharose excel resin and the protein of interest was collected in the unbound fraction. Protein samples were transferred into 50 mM potassium phosphate buffer pH 7.5 using HiPrep 26/10 desalting column. All protein samples were analyzed by SDS-PAGE gel electrophoresis and the protein concentration was determined based on absorbance and extinction coefficients calculated from the amino acid sequence. Correct protein identity was confirmed by matrix-assisted laser desorption/ionization time-of-flight mass spectrometry (MALDI-TOF/MS).

### Phage display and analysis of NGS results

The RiboVD phage library displays the IDRs of mammalian and avian RNA viruses (Riboviria; taxonomic identifier: 2559587) tiled aby 16 amino acids overlapping peptides (**Table S1**) (Kruse et al., 2021). The library design is available on-line (http://slim.icr.ac.uk/phage_libraries/rna_viruses/species.html). The library was used in triplicate phage selections against 139 His-GST/MBP tagged bait protein domains (Table S2). In brief, proteins (10 µg in 100 µL PBS) were immobilized in 96 well Flat-bottom Immunosorp MaxiSorp plates for 18 h at 4°C. Wells were blocked with 200 µL BSA (0.5% in PBS) and washed four times with 200 µL PT (PBS+0.05% (v/v) Tween20) before adding the phage library (10^11^ phage in 100 µL PBS per well), first to the GST-coated wells (1 h) to remove non-specific binders, and then to the bait protein-coated plates (2 h). Unbound phages were removed and the bound phages were eluted (100 µL log phase *E. coli* OmniMAX, 30 min, 37°C). M13 helper phages were added (10^9^ M13KO7 helper phages per well, 45 min at 37°C) before transferring the bacteria to 1 mL 2xYT supplemented with 100 µg carbenicillin (Carb), 30 µg Kan and 0.3 mM IPTG. Bacteria were grown at 37°C for 18 h, before harvesting the phages (2,000 x g for 10 min). The phage supernatants were pH adjusted (using 1/10 volume 10x PBS) and used as in-phage for the next round of selection.

The peptide-coding regions of the naive RiboVD library and the binding-enriched phage pools (5 µL) were PCR-amplified and barcoded using Phusion High-Fidelity polymerase (Thermo Scientific) for 22 cycles. PCR products were confirmed by 2% agarose gel electrophoresis stained with GelRed using a 50 bp marker (BioRad). PCR products were normalized using Mag-bind Total Pure NGS, pooled and purified from a 2% agarose gel (QIAquick Gel extraction Kit), and analyzed using Illumina MiSeq v3 (1x150 bp read setup, 20% PhiX). Results were processed using in-house Python scripts. Reads were demultiplexed, adapter and barcode regions were trimmed, and sequences were translated into peptide sequences. Peptides were annotated using PepTools (Benz et al., 2022) and assigned confidence levels based on four different criteria: occurrence in replicate selections, identification of overlapping peptide sequences, high counts, occurrence of sequences matching consensus motifs determined from the generated data set, or a priori defined consensus motifs for the bait proteins (Benz et al., 2022; Kumar et al., 2022). For a stringent analysis we focused on the medium/high confidence peptides that fulfill at least three of these criteria. The state of viral protein annotation in UniProt is ever-changing, and multiple strains of the same viral species sometimes have multiple entries for the same (or very similar) proteins and polyproteins. Our annotations with PepTools takes this situation into account. When counting the number of interactions, we opted to collapse the viral proteins based on a combined IDs that include their names (not accessions), chain names (not chain IDs), and species (at the species level, not strain).

### Viral network generation and analysis

The Human PPI network was extracted from IntAct (version: 4.2.17, last update May 2021) (Orchard et al., 2014). We also included kinase-kinase interactions and kinase-substrate interactions from PhosphoSitePlus (Orchard et al., 2014) (version 6.5.9.3, last update May 2021), OmniPath (Turei et al., 2016) (last release May 2021) and SIGNOR 2.0 (Licata et al., 2020) (last release May 2021). Only proteins annotated in Swiss-Prot (UniProt, 2021) and those annotated with at least one GO term (Gene Ontology, 2021) were kept. The resulting protein interaction network (PIN) comprises 16,407 nodes and 238,035 edges. Edge weights are modeled according to the Topological Clustering Semantic Similarity (Jain and Bader, 2010) and calculated using the Semantic Measure Library (Harispe et al., 2014). Additionally, to determine the significance associated with each node, we generated 1000 random networks employing the configuration model available in the python igraph library (http://igraph.org) updating the edge weight accordingly. Each network is Laplacian-normalized to correct for the hub bias. In the formula:

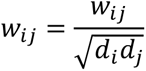

where *w_ij_* indicates the edge weight (i.e. semantic similarity) and *d_i_* and *d_j_* represent the weighted degree of node *I* and node *j*.

The Random walk with restart (RWR) algorithm (though the personalized PageRank function available in http://igraph.org was used to simulate the propagation of viral infection into the PIN. The human proteins targeted by the virus were selected as seed nodes for the RWR procedure selecting a restart probability equal to 0.7. The RWR algorithm was also executed on the 1000 random networks employing the same seed nodes and restart probability. This allows us to estimate the empirical p-value for each protein in the PIN as the percent of random score that exceeded the real score (excluding the seed genes), that is:

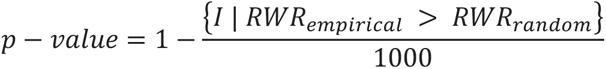

Where I is the indicator function, *RWR_empirical_* and *RWR_random_* refer to the RWR score assigned to the empirical PIN and the random networks respectively. Only nodes with a *p* −*value*< 0.01 are considered significant. In total 575 target networks, one for each virus, from 26 different viral families are extracted. Each target network is represented by a vector comprising the significant RWR scores associated with the proteins belonging to the target network. To identify the common biological processes subjected to the viral interference, the human nodes in the networks that are significantly affected by viral infection are selected. To do so, for each protein in the networks, we defined the average RWR family specific score as:

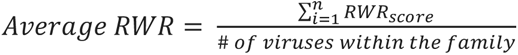

Representing the average RWR score assigned to each significant protein belonging to the respective family. To assess which average RWR score is significant, we calculated the upper-tailed Z-Score test, employing as background distribution the random walk scores of those nodes that didn’t pass the significance threshold (i.e. *p* − *value* > 0.01). Proteins with a score in at least 8 viral families and with a Z-Score > 2.32 (corresponding to a *p* − *value* < 0.01) were selected. This set constitutes the foreground for the enrichment analysis against GO using the human proteome as background. We used g:Profiler (Raudvere et al., 2019) to perform enrichment analysis (Table S6), focusing on the biological process domain. Then, we employed Enrichment Map (Merico et al., 2010) and Cytoscape (Shannon et al., 2003) to visualize the GO biological process map.

### Cluster network families

Firstly, for each of the 575 viral signatures, we performed an enrichment analysis against Reactome (Gillespie et al., 2022) using all levels of the pathway hierarchy. Fisher’s exact test (Fisher, 1934) based on the hypergeometric distribution is used to determine the overrepresented terms and the Bonferroni correction (Bonferroni, 1936) is applied to correct for multiple comparisons. To extract the 26 family signatures, we summed the corresponding RWR scores of the proteins in the viral signature vectors appertaining to their respective family. After this procedure, a matrix **A** of 26 X 4275 elements is obtained, where each row corresponds to a family signature and each column represents the sum of the RWR scores for each protein within their respective family. A value equal to 0 is assigned if the protein was not significant in any of the viral signatures within that family. Since the distribution of the viruses inside each viral family was different, we normalized the matrix using the quantile normalization from the *scikit-learn* package (Pedregosa et al., 2011). Next, since the normalized matrix was positive, we applied the standard non-negative matrix factorization (NMF) from the *nimfa* library with default parameters (latent factor a part) (Zitnik and Zupan, 2012) to identify groups of viral families targeting similar human pathways. A critical step in NMF was to select the right number of latent factors. For this aim, we ran the NMF algorithm 1000 times employing the initialization algorithm to obtain a stable consensus clustering (Lee and Seung, 1999). In each run, we calculated the cophenetic correlation coefficient. We selected 5 latent factors as evident from the violin plot (Figure S3), since increasing the number of latent factors slightly increased the cophenetic correlation coefficient. Hence the normalized matrix **A** was decomposed into:

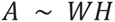

The maximum value on each row of the coefficient matrix **H** represents the strongest membership of the family with the latent component and consequently a cluster. We calculated mean of the relative frequency of a Reactome pathway within the family inside each cluster:

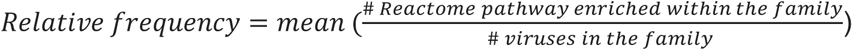

and the absolute frequency of that pathways inside the cluster:

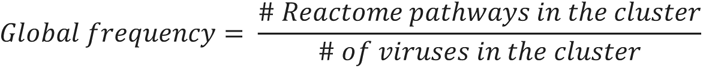

To consider a Reactome pathway representative of each cluster both scores must be greater than 0.2 (see Figure 2B).

To compare the vesicle-mediated transport networks, we extracted all the enriched proteins involved in the endocytosis pathway for cluster 4 and 5 respectively for each of the families involved and analyzed them using Cytoscape.

### Affinity measurements

Affinity measurements were performed in 50 mM potassium phosphate, pH 7.5, or 50 mM potassium phosphate, pH 7.5, 1 mM TCEP. Experimental setup and conditions were identical for all domains unless stated otherwise. The affinity between the protein domains and their respective FITC-labeled peptides was determined with saturation binding experiments (Figure S5). A 1:1 dilution series with increasing concentration of protein of interest was performed containing a fixed concentration of FITC-labeled peptide (ranging from 5-10 nM depending on the protein under investigation) in black, non-binding surface, flat bottom 96-well plates. Measurements were performed on a SpectraMax iD5 plate reader at room temperature and at excitation/emission wavelengths of 485/535 nm. The G-factor was set accordingly so that the wells containing only the FITC-labeled peptide showed a fluorescence polarization value between 10-40 mP (corresponding to B_bottom_). Saturation binding curves were analyzed by GraphPad Prism and fitted to the equation

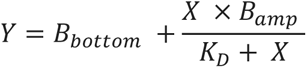

where B_bottom_ is the fluorescence polarization value of FITC-labeled peptide in absence of protein, B_amp_ is the amplitude of fluorescence polarization signal (B_top_ - B_bottom_), X is the concentration of free protein (equal to total protein since [protein]>>[FITC-peptide]), *K*_D_ is the equilibrium dissociation constant and Y is the fluorescence polarization signal.

To determine affinities between proteins and non-labeled peptides a competition assay was performed. The non-labeled peptide was added at increasing concentrations to a fixed concentration of FITC-labeled peptide (5-10 nM final concentration, depending on the protein) and protein of interest. Fixed concentrations of proteins in displacement experiments were as follows to achieve approximately 60% saturation of the complex between protein and labeled peptide: ALIX V: 4-6 µM, TSG101 UEV: 8 µM, NEDD4 WW2: 30 µM, NEDD4 WW4: 30 µM, CEP55 EABR: 1 µM, GGA3 VHS: 15-17 µM, GGA3 ear: 4 µM, CLTC NTD: 30 µM, AP2M1: 0.9-1.65 µM and PABP1 C: 1.76-2 µM. FP values from the competition assay were fitted (GraphPad Prism) to a sigmoidal dose-response equation

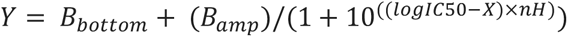

where Y is the fluorescence polarization signal, B_bottom_ is the FP value of FITC-labeled peptide in absence of protein, B_amp_ is the amplitude of FP signal (B_top_ - B_bottom_), IC50 is non-labeled peptide concentration required for 50% apparent inhibition, X is the logarithmic value of non-labeled peptide concentration and nH is the Hill coefficient. The resulting IC50 values obtained from the displacement experiment were converted to *K*_D_ values as previously described (Nikolovska-Coleska et al., 2004). All *K*_D_ values were calculated on the raw fluorescence polarization data. Normalization was employed to facilitate easier visualization. All saturation and competition experiments were performed at least in three technical replicates.

### Crystallization

The CLTC NTD was co-crystallized with two viral peptides that were also used in affinity measurement studies namely Nsp3_1765-1780_ (EEEV) and mu-NS_705-720_ (MRV1), by vapor diffusion method (MRC 2 Well Crystallization Plate in UVXPO; Hampton research). CLTC NTD concentrated to 18 mg/ml in 50 mM Tris-Cl (pH-7.7), 200 mM NaCl, 4 mM DTT was mixed with peptides dissolved in the same buffer at 10 mg/ml at a protein:peptide ratio of 1:2 and stored at -20°C until crystallization plate setup. Initially, the crystallization was attempted by using reported crystallization conditions (50 mM Tris-Cl pH-7.5 and 30 % PEG 6000) (Rondelet et al., 2020). The crystal growth was optimized by varying the pH of Tris (pH 7.0 – 8.5) and concentration of PEG 6000 (20-30%). For both peptides the plate-like crystals appeared in several drops within 2 days. Microseed stocks were prepared for each of the CLTC NTD- peptide complexes from the crushed crystals harvested from a single drop, diluted 1:100 with the respective mother liquors. These stocks were used to screen the conditions of the Morpheus crystallization screen (Gorrec, 2015) in a sitting-drop setup. For each complex, single crystals appeared under several conditions. The best diffracting CLTC NTD-Nsp3_1765-1780_ (EEEV) crystals were grown using 30% PEG 550 MME/PEG 20K and 0.1 M NPS buffer system pH 6.5 (containing NaNO_3_, Na_2_HPO_4_ and (NH_4_)_2_SO_4_) as reservoir solution. The best CLTC NTD-mu-NS_705-720_ (MRV1) crystals were obtained with 30% PEG 550 MME/PEG 20K, 0.12 M monosaccharides (D-Glucose, D-Mannose, D-Galactose, L-Fucose, D-Xylose, N-Acetyl-D-Glucosamine) and 0.1 M sodium HEPES/MOPS pH 7.5. Crystals were cryo-cooled in liquid nitrogen without additional cryoprotectant.

PAPB1 PABC domain was concentrated to 20 mg/ml in 50 mM Tris (pH-7.5), 150 mM NaCl, 1 mM DTT and incubated with the N_351-366_ (HCoV229E) peptide at 1:1.5 molar ratio. The ammonium sulfate screen (AmSO_4_ suit, Hampton Research) was used to identify the initial crystallization conditions at 22°C. The crystallographic data were collected from crystals grown using a reservoir solution of 0.1 M sodium MES pH 6.5, 1.8 M ammonium sulfate. Crystals were briefly soaked in mother liquor containing 20% glycerol prior to cryo-cooling in liquid nitrogen.

### X-ray data collection, structure determination and refinement

For the two peptide complexes of CLTC, crystallographic data was collected at 100 K at the beamline I04 of the Diamond Light Source (Didcot, UK) and processed on site using either Fastdp or Xia2 (Winter and McAuley, 2011). The structures were solved by molecular replacement using Phaser (McCoy et al., 2007) and PDB entry 1C9I as search model (ter Haar et al., 2000). The PABPC1 PABC-HCoV 229E data were collected at BioMAX, MAX IV (Ursby et al., 2020) (Lund, Sweden), and processed at the beamline using the autoproc pipeline (Vonrhein et al., 2011). The structure was solved by a molecular replacement method using Phaser and PDB entry 3KUJ (Kozlov and Gehring, 2010) as the search model. All three structures were refined with phenix.refine and Refmac5 of the Phenix (Adams et al., 2010) and CCP4 program suites (Winn et al., 2011), respectively. Manual model building was done in *Coot* (Emsley and Cowtan, 2004). The final structures showed good geometry as analyzed by Molprobity (Chen et al., 2010). The data collection and refinement statistics are given in Table S9.

### Cells and viruses

Human embryonic kidney 293 cells (HEK293) (Sigma), HEK293T (TakaraBio), and African green monkey kidney E6 cells (VeroE6) cells (ATCC, CRL-1586) were cultured in Dulbecco’s modified Eagle’s medium (DMEM)(Gibco) supplemented with 10% (v/v) fetal bovine serum (FBS) (HyClone) and 100 units/ml penicillin G with 100 μg/ml streptomycin solution (PEST) (Gibco) at 37°C, 5% CO_2_, humidified chamber unless otherwise specified. The African green monkey kidney B4 cells (VeroB4) cells were cultured in 199/EBSS medium (HyClone) supplemented with 10 % (v/v) FBS, and PEST. For PLA, HEK293 and HEK293 overexpressing HA-tagged human PDGFRβ (HEK293-PDGFRβ-HA, a kind gift from Frank Böhmer (Markova et al., 2003; Tenev et al., 2000)) were cultured in DMEM and Nutrient Mixture F-12 (1:1) (Gibco) supplemented with 10% (v/v) FBS (Gibco) and PEST.

SARS-CoV-2 (SARS-CoV-2/01/human2020/SWE accession no/GeneBank no MT093571.1, provided by the Public Health Agency of Sweden), was grown in VeroE6 cells and used at passage number 4. Japanese encephalitis virus (JEV) (Nakayama strain), West Nile virus (WNV) (WNV_0304h_ISR00), yellow fever virus (YFV) (Asibi), and dengue virus (DENV) (serotype-2; PNG/New Guinea C) were kind gifts from S. Vene, the Public Health Agency of Sweden and were grown in VeroB4 cells. TBEV (Torö-2003, (Asghar et al., 2016), Langat virus (LGTV) (TP21, kind gift from Gerhard Dobler Bundeswehr Institute of Microbiology, Munich, Germany), ZIKV (MR766, kind gift from Gerhard Dobler Bundeswehr Institute of Microbiology, Munich, Germany), RVFV (Islam et al., 2018), vesicular stomatitis viruse (VSV) (kind gift of Friedemann Weber, University of Freiburg), Sindbis virus (SINV) (Lovanger, KF737350, kind gift from Olivia Wesula Luande) and chikungunya virus (CHIKV) (CHIKV LR2006OPY1, kind gift from Magnus Evander) were grown in VeroB4 cells.

### GST-pull down assay

The GST pulldown assay was performed as described previously (Inturi et al., 2013). Whole cell lysates were obtained by transfecting HEK293T cells cultured on 100 mm culture plates with plasmids expressing C-terminal Flag-tagged NP (ZEBOV) wt, NP (ZEBOV) mut 1, NP (ZEBOV) mut 2, Nsp3 (EEEV) wt, Nsp3 (EEEV) mut, N (HCoV 229E) wt, N (HCoV 229E) mut, P (HeV) wt and P(HeV) mut proteins. 48 hours post transfection, the cells were washed with 1 X PBS and lysed in GST-lysis buffer containing 25 mM Hepes-KOH (pH 7.4), 12.5 mM MgCl_2_, 100 mM KCl, 0.1 mM EDTA, 10% glycerol, 0.1% NP-40, supplemented with protease inhibitor for 30 minutes on ice. The cell lysates were freeze-thawed three times and the supernatant was collected by centrifugation at maximum speed for 15 min. The cell lysates were incubated with GST-tagged proteins for 1 hour, at room temperature with end-over-end mixing. The beads were washed with the GST-lysis buffer and the bound proteins were separated by SDS-PAGE and analyzed by western blotting. For western blotting, the SDS-PAGE separated proteins were transferred onto nitrocellulose membrane (Amersham, Protran) for 2 hours, 200 mA at 4°C. The membrane was blocked in Odyssey blocking buffer (LI-COR) for 1 hour at room temperature and incubated in primary antibodies anti-mouse Flag (Sigma, M2, F1804), anti-rabbit GST (Santa Cruz, sc-33614), overnight at 4°C. The membrane was washed three times in PBS-T (PBS+0.1% Tween 20) before incubation with fluorescent secondary antibodies (IRDye®, LI-COR) against anti-mouse or anti-rabbit for 30 min at room temperature. The membrane was washed three times in PBS-T and scanned using Odyssey scanner (LI-COR).

### Proximity ligation assays

HEK293 cells were seeded in 8-well Nunc Lab-Tek II chamber slides (0,7 cm^2^, Sigma) at a density of 70.000 cells/cm^2^. After 40 hours, the cells were transfected with plasmids expressing C-terminal Flag-tagged Nsp3 (EEEV) wt, Nsp3 (EEEV) mut, P (HeV) mut proteins (Table S9) or not transfected. Growth medium was replaced with Opti-mem (ThermoFisher) and the cells transfected with 100 ng Plasmid DNA per well using Lipofectamine 3000 (ThermoFisher) as described by the manufacturer. After 6 hours of incubation, the medium was replaced with the growth medium and grown overnight. On ice, cells were washed in ice cold PBS, then fixated in ice cold formalin solution (3.7% paraformaldehyde plus 1% methanol in PBS) for 15 minutes before washing in PBS 3 times for 5 minutes. The slides were dried and the wells encircled with an ImmEdge hydrophobic barrier pen (Vector Laboratories). The slides were rehydrated in TBS and the cells permeabilized in TBS plus 0.2% Triton X-100 for 10 minutes. In a moisture chamber, the slides were blocked in blocking buffer consisting of Odyssey Intercept (TBS) Blocking Buffer (Licor) plus TBS in a 1:1 ratio for 1 hour at 37°C, before incubation overnight at 4°C with primary antibodies goat-anti-FLAGtag (ab1257, abcam) (1:1000) and mouse-anti-clathrin (ab2731, abcam) (1:200) diluted in blocking buffer. The slides were washed 3 times 10 minutes in TBS plus 0,05% Tween-20 before incubation with Duolink secondary probes (Olink) compatible with host species of the primary antibodies. The slides were incubated for 1 hour at 37°C with Duolink PLA probe anti-Mouse PLUS and Duolink PLA probe anti-Goat MINUS diluted in blocking buffer to a concentration of 1x. The slides were washed 3 times for 10 minutes in TBS plus 0,05% Tween-20 and incubated with 1x Duolink Ligation solution and 1 U/μL T4 DNA ligase (Thermo Fisher) for 30 minutes at 37°C. The slides were washed 3 times for 10 minutes in TBS and incubated with 1x Duolink Amplification Red solution and 0.125 U/μL Phi 29 polymerase (Montserate) and washed again 3 times for 10 minutes in TBS. To visualize transfected cells, the slides were incubated with Donkey anti-goat Alexa Flour Plus 647 (A32849, Thermo Fisher) and Hoechst 33342 for 1 hour at 37°C. The slides were washed again 3 times for 10 minutes in TBS plus 0,05% Tween-20, then briefly washed in TBS and mounted with Slowfade Gold antifade mounting reagent (S36936, Thermo Scientific).

PLA experiments with PDGFRβ were performed using HEK293 overexpressing HA-tagged human PDGFRβ (HEK293-PDGFRβ-HA). The PLA experiments were performed as described above, except after transfection the cells were starved overnight in starvation medium (DMEM/F-12, 0,2%FBS) and then stimulated with 50 ng/ml PDGF-BB (Peprotech) in starvation medium for 0, 10 and 60 minutes at 37°C before fixation. Primary antibodies used were rabbit-anti-PDGFRβ (#3169, Cell Signaling Technology) (1:100) and mouse-anti- PDGFRβ-pY751 (#3166, Cell Signaling Technology) (1:200), and Duolink PLA probes were anti-Mouse PLUS and Duolink PLA probe anti-Rabbit MINUS. To visualize transfected cells, the slides were incubated with FLAG-tag antibody (1:1000) for 1 hour at room temperature, washed 3 times for 10 minutes in TBS plus 0,05% Tween-20, and subsequently incubated with secondary antibody Donkey anti-goat Alexa Fluor Plus 647 (A32849, Thermo Fisher) diluted 1:500 and 10 μg/mL Hoechst 33342 in blocking buffer for 1 hour at 37°C. The slides were washed 3 times for 10 minutes in TBS plus 0,05% Tween-20 and mounted as previously described.

Slides were imaged using a Zeiss Imager Z2 controlled by Zen 2 (blue edition) software. The microscope was equipped with a Hamamatsu C11440 camera, a 40x/1.4 oil objective, filter cube sets 31, 43 HE, 49, and 50 from Zeiss, and a HXP 120V light source set to 90% for all channels imaged. 3 images per condition for each experiment were acquired as z-stacks of 11 slices 0.5 μm apart. The images shown are the maximum intensity projection of the z-stack and have been adjusted for brightness and contrast for visualization purposes.

Image analysis and quantification of PLA signal was performed using CellProfiler software version 3.0.0 or newer, made available by the Broad Institute Imaging Platform (McQuin et al., 2018). Image analysis was performed on the maximum intensity projection of the z-stack of original images. Segmentation of the cells was performed based on the image resulting from the Hoechst channel using first the IdentifyPrimaryObjects module for segmentation of nuclei based on a global three-class Otsu threshold method using intensity to distinguish and draw dividing lines between clumped objects, followed by the IdentifySecondaryObjects module to segment cells using the Distance-N function with a fixed maximum distance from the nucleus to cell border. The PLA signal was evaluated as PLA rolling circle amplification product (RCP) per cell. The image from the TexasRed channel was first filtered with the help of the EnhanceOrSuppress module to enhance the feature type “speckles” and remove background. The filtered image from the TexasRed channel was then used as input for segmentation of RCPs, based on manual thresholding using the IdentifyPrimaryObjects module. RCPs were then related to the cells via the RelateObjects module. Integrated intensity per cell was measured using the MeasureObjectIntensity module for the channel imaging the FLAGtag. Finally, all intensity measures and RCPs per cell were exported to an Excel spreadsheet. To distinguish data from transfected and nontransfected cells, a cutoff intensity for transfected cells was set corresponding to the highest integrated intensity per cell of the FLAGtag containing channel for nontransfected cells.

### Cell surface fluorescence assay

HEK293-PDGFRβ-HA cells were seeded, transfected and stimulated for 0 or 60 minutes as described for PLA experiments. On ice, the cells were washed in ice cold PBS and incubated with a primary antibody targeting the extracellular part of PDGFRβ, 5 μg/ml goat-anti-PDGFRβ (AF385, RnD Systems) in PBS for 1 hour. The cells were washed 3 times for 10 minutes in PBS before fixation, permeabilization and blocking was performed as described for PLA experiments. The cells were incubated with rabbit-anti-FLAG (1:800) (#14793S, Cell Signaling Technology) diluted in blocking buffer overnight at 4°C, washed 3 times 10 minutes in TBS plus 0,05% Tween-20, and subsequently incubated with secondary antibodies Donkey anti-rabbit Alexa Fluor Plus 555 (A32794 ThermoFischer) and Donkey anti-goat Alexa Fluor Plus 647 (A32849, ThermoFischer) diluted 1:500 and 10 μg/mL Hoechst 33342 Solution (Thermo Scientific) (1:1000) in blocking buffer. The slides were washed 3 times for 10 minutes in TBS plus 0,05% Tween-20 and mounted and observed under microscope as described for PLA experiments. Images were analyzed with CellProfiler, using the same pipeline for segmentation and distinguishing between transfected and nontransfected cells as described for PLA experiments. Fluorescence intensity was measured as integrated intensity per cell for the channel imaging PDGFRβ using the MeasureObjectIntensity module.

### Lentivirus plasmids and production

Lentiviruses were produced by transfection of HEK293T cells in 100 mm plates as described previously (Kruse et al., 2021). To produce lentiviruses, pLJM1-EGFP (David Sabatini lab, Addgene plasmid #19319; (Sancak et al., 2008)), psPAX2 (Didier Trono lab, Addgene plasmid #12260), and pMD2.G (Didier Trono lab, Addgene plasmid #12259) were used. To generate pLJM1-EGFP transfer plasmids, four copies of inhibitory peptide or control peptide with mutated binding motif spaced out by a flexible GST linker and fused to C-terminus of EGFP were ordered from GenScript. At 72 h post transfection, the supernatants from cells transfected with lentivirus plasmids were filtered and stored at -80°C.

### Viral infections

VeroE6 or VeroB4 cells were seeded into greiner CELLSTAR® 96-well plates containing EGFP-PABPi mut or EGFP-PABPi lentivirus (**Figure 6F**) in DMEM containing 2 % FBS and 1 μg/mL polybrene, and incubated for 72 h. Transduced cells were then infected with a panel of RNA viruses (VeroE6: SARS-2 (MOI: 0.05 for 16 h), JEV (MOI: 0.1 24 h), WNV (MOI: 0.1 24 h), YFV (MOI: 0.1 24 h), ZIKV (MOI: 0.1 24 h), RVFV (MOI: 0.05 for 16 h), VSV (MOI: 0.001 5 h), SINV (MOI: 0.05 for 16 h) and CHIKV (MOI: 0.05 for 16 h), VeroB4: DENV, TBEV and LGTV with MOI:0.1 for 24 h. Virus was detected using the following primary antibodies, SARS-2 (SARS-CoV-2 nucleocapsid (Rabbit monoclonal, Sino Biological Inc., 40143-R001)), JEV, WNV, DENV and ZIKV (mouse monoclonal anti-flavivirus E HB112 ATCC), YFV (YFV E CRC 1689 ATCC), TBEV and LGTV (mouse monoclonal anti-TBEV E 1786, PMID: 7817895[RL1]), VSV, SINV and CHIKV (mouse monoclonal to J2 (Scicons 10010500)), and secondary antibodies either donkey anti-mouse or donkey anti-rabbit IgG Alexa Fluor 555 secondary antibody (Invitrogen). Nuclei were counterstained by DAPI. Number of infected cells were determined using a TROPHOS Plate RUNNER HD® (Dioscure, Marseille, France). Number of infected cells were normalized to DAPI count and presented as percentage infection of mutated peptide.

### Viral titrations

SARS-CoV-2 was diluted in ten-fold dilutions and added to VeroE6 cells followed by 1 h incubation. The inoculum was replaced with an overlay containing DMEM, 2% FBS, 1% PEST and 1.2% Avicel. After 24 h of infection cells were fixed in 4% formaldehyde for 30 minutes, permeabilized in PBS 0.5 % trition-X-100 and 20 mM glycine. Viral foci were detected using primary monoclonal rabbit antibodies directed against SARS-CoV-2 nucleocapsid (Sino Biological Inc., 40143-R001), and secondary anti-rabbit HRP conjugated antibodies (1:2000, Thermo Fisher Scientific). Viral foci were then revealed by incubation with TrueBlue peroxidase substrate for 30 minutes (KPL, Gaithersburg, MD). TBEV was titrated as previously described (PMID: 29544502).

### Immunofluorescence microscopy of EGFP-PABPi transfected cells

VeroB4 cells expressing either EGFP-PABPi mut or EGFP-PABP inhibitor peptides (EGFP-PABPi) were seeded in 8-well chamber slides (Sarstedt) and infected with TBEV at an MOI of 1 for 24 hours. The cells were fixed with 4% formaldehyde and incubated with permeabilization buffer (0.3% TritonX-100 and 1% Goat serum in PBS) containing primary antibodies against dsRNA J2 ((1:1000) Scicons 10010500) and PABPC1 ((1:100) Abcam ab21060) followed by incubation with DAPI (1:1000) and conjugated secondary antibodies anti-mouse Alexa555 and anti-rabbit Alexa647 (1:500, Thermo Fisher Scientific). Coverslips were mounted and samples were analyzed using a Leica SP8 Laser Scanning Confocal Microscope with a 63x oil objective (Leica) and Leica Application Suit X software (LAS X, Leica). For the quantification of the RadialCV a total of 6 images containing 185 and 158 infected cells from EGFP-PABPi mut and EGFP-PABPi, respectively, were analyzed using CellProfiler. The DAPI channel was used to identify the nuclei as primary objects while the PABPC1 channel was used to identify the whole cells as secondary objects. These two objects where then used to identify the cytoplasmic fraction as a tertiary object. The cytoplasmic fraction was analyzed with the “MeasureObjectIntensity” and “MeasureObjectIntensityDistribution” functions to determine infected cells using the dsRNA integrated intensity and create the fractions within the cytoplasm to determine the distribution of dsRNA signal using the RadialCV.

### AP-MS

The growth media contained 10% FBS (Gibco), non-essential amino acids (NEAA, Gibco) and 5 µg/mL and 5 units/mL penicillin-streptomycin (Gibco). One T175 flask of HEK293 cells of 70% confluency per condition was transiently transfected using 90 µg of EGFP-PABPi or EGFP-PABPi mut, and Lipofectamine 3000 (Invitrogen) according to manufacturer’s instructions. The cells were harvested 24 hours after transfection by first washing with ice cold DPBS (Gibco) then scraped into 3 ml ice cold lysis buffer (10 mM Tris-HCl, pH 7.5, 150 mM NaCl, 1% NP-40 substitute (Sigma 74385), 1x Protease inhibitor (Roche, cOmplete, Mini, EDTA-free, 4693159001) and incubated on ice for 30 min while shaking. The lysate was clarified by centrifugation at 16000 g for 15 minutes at 4°C. Similarly prepared, but SARS-CoV-2 or TBEV infected VeroE6 or VeroB4 cells, respectively, were also used, stably expressing the above-mentioned constructs. The protein concentration was determined using DC Protein Assay (Bio-Rad).

The cell lysate was diluted to 0.8 mg protein/ml with dilution buffer (10 mM Tris-HCl, pH 7.5, 150 mM NaCl, 1x Protease inhibitors), and 1 mg protein was used per replicate. Cell lysates were incubated with GFP-Trap® Dynabeads™ (Chromotek) at 4°C for 1 hour while rotated. After washing, the interacting proteins were eluted using acidic elution buffer (200 mM glycine-HCl, pH 2.5) and neutralized with 1 M ammonium bicarbonate instantly. The eluate was reduced with DTT and alkylated with IAA, then digested overnight using trypsin at 37°C. The digestion was stopped using an acidifying solution (83.3% AcN, 16.7% TFA) to pH<3. The peptides were desalted using STageTips made in-house (Rappsilber et al., 2003; Rappsilber et al., 2007), with centrifugal elution. Briefly, 2 layers of C18 membrane (3M Empore) were placed in a 200 µl pipette tip, activated with methanol and 80% AcN, 0.1% formic acid, then washed twice with 0.1% formic acid. After that the acidified samples were loaded, washed with 0.1% formic acid and eluted with 80% AcN, 0.1% formic acid. The eluted sample was vacuum-dried and stored at -80°C.

The samples were analyzed using an Easy-nLC 1000 nanoLC (Thermo) with an Acclaim PepMap 100 pre-column (Thermo, 75 µm x 2 cm, 3 µm, 100Å) and a PepMap RSLC C18 analytical column (Thermo, EASYspray, 75 µm x 15 cm, 2 µm, 100Å). The mass spectrometer was a QExactive Plus Orbitrap instrument (Thermo) equipped with an EASYspray ion source. For peptide separation, a gradient method was applied, where the gradient went from 4 to 76% acetonitrile in 79 minutes. The MS was operated in the positive ion mode with a resolution of 140000 for full scan (400-1700 m/z), and 17500 for MS/MS with the automatic gain control (AGC) target of 3x10^6^ and 1x10^5^, respectively. The ESI spray voltage was 1.9 kV. Data-dependent acquisition was used, with the top 10 most abundant ions fragmented and measured in MS/MS. Dynamic exclusion of 30 seconds was enabled.

The raw files were analyzed using MaxQuant (version 2.0.1.0) using FASTA files acquired from Uniprot: *Homo sapiens* (2022.02.21, reviewed, 20360 entries) for HEK293 samples and *Chlorocebus* (2022.02.22, reviewed and unreviewed, 20717 entries) for VeroE6 samples with or without proteins of the SARS-CoV-2 variant patient isolate SARS-CoV-2/01/human/2020/SWE accession no/GeneBank no MT093571.1. or Tick-borne encephalitis virus Torö-2003, GenBank Accession no. DQ401140.3. Trypsin/P was selected as the digestion enzyme, with maximum 2 missed cleavages allowed. For variable modifications methionine oxidation and N-terminal acetylation were allowed, while for fixed modification carbamidomethylation of cysteines was selected. Label-free quantification was chosen using the MaxLFQ algorithm (Cox et al., 2014) and a minimum ratio count of two. The used peptide mass tolerances were 20 and 4.5 ppm for first and main search, respectively. PSM and protein FDR was set to 0.01. The minimum number of detected peptides was set to 2, and the minimum number of unique peptides to 1 for identification.

To identify interacting proteins, the data was processed first with Perseus (2.0.3.0) (Tyanova et al., 2016). Using the proteingroups.txt result file from MaxQuant, the possible contaminants, reverse hits and proteins only identified by site were removed. The LFQ intensities were transformed to a log_2_(x) base, and the hits were filtered, only keeping rows with at least 3 valid values in at least one of the categorical groups (sample/control). The missing values were replaced from normal distribution with a width of 0.3 and down shift of 1.8 (mode: total matrix). Two-sided t-test was used for significance testing (p-value <0.05, S0:0) and the results were visualized in a Volcano plot using a fold-change cut off of 2.

## Supporting information

Tabel S7

## ACKNOWLEDGEMENTS

This work was supported by the grants from the Swedish Foundation for Strategic research (YI, PJ: SB16-0039), the Swedish Research Council (YI: 2020-03380; PJ: 2020-04395; AKÖ: 2018-05851), the Knut and Alice Wallenberg Foundation (YI, PJ, and AKÖ via Science for Life Laboratory, 2020.0182) and a Cancer Research UK Senior Cancer Research Fellowship (ND: C68484/A28159). GG and EP were funded by the European Molecular Biology Laboratory. We thank the medical faculty Umeå University strategic research resource, the Laboratory for Molecular Infection Medicine Sweden (MIMS), and Umeå Center for Microbial Research (UCMR) for generous support (A.Ö.), and the Biochemical Imaging Center at Umeå University and the National Microscopy Infrastructure, NMI (VR-RFI 2016-00968) for assistance in microscopy. Sequencing was performed by the SNP&SEQ Technology Platform in Stockholm. The facility is part of the National Genomic Infrastructure (NGI) Sweden and Science for Life Laboratory and is also supported by the Swedish Research Council and the Knut and Alice Wallenberg Foundation. Work at the Novo Nordisk Foundation Center for Protein Research is supported by grant NNF14CC0001.

## AUTHOR CONTRIBUTION

Conceptualization: OS, PJ, NED, YI; Investigation: FM, LS, GG, MRS, RL, MBAP, CB, EK, DB, RI, MA, AS, EA, HA; Data curation: LS, IK, NED; Writing - original draft: FM, MRS, EP, NED, PJ, YI; Writing - review & editing: LS, AKÖ, RI, EK, OS; Visualization: FM, LS, NED, MRS, MBAP, RI; Supervision: YI, OS, DD, EP, PJ, AKÖ, NED; Project administration: YI; Funding acquisition: YI, PJ, OS.

## DATA AVAILABILITY

The mass spectrometry proteomics data have been deposited to the ProteomeXchange Consortium via the PRIDE (Perez-Riverol *et al.,* (2022)) partner repository with the dataset identifier PXD033874. The crystal structures have been deposited in PDB and are available with the PDBis 7BN1, 7BN2 and 7BN3. The interaction data will be deposited to IntAct. Relevant code has been made available (https://zenodo.org/deposit/6583610). All datasets will be made available upon publication.

## SUPPLEMENTAL TABLES (see separate files)

**Table S1.** RiboVD library design.

**Table S2.** Domain collection.

**Table S3.** RiboVD selection data.

**Table S4.** RiboVD motif benchmarking set.

**Table S5.** RiboVD PPI benchmarking set.

**Table S6.** GO term enrichment of expanded hosts-virus network.

**Table S7.** Compiled FP affinity measurements data and peptide information.

**Table S8.** Full length viral proteins and lentiviral constructs used in cell base experiments.

**Table S9.** Crystallographic data.

**Table S10.** AP-MS data.

**Table S11.** Reagents and resources.

## SUPPLEMENTAL FIGURES

**Figure S1.**
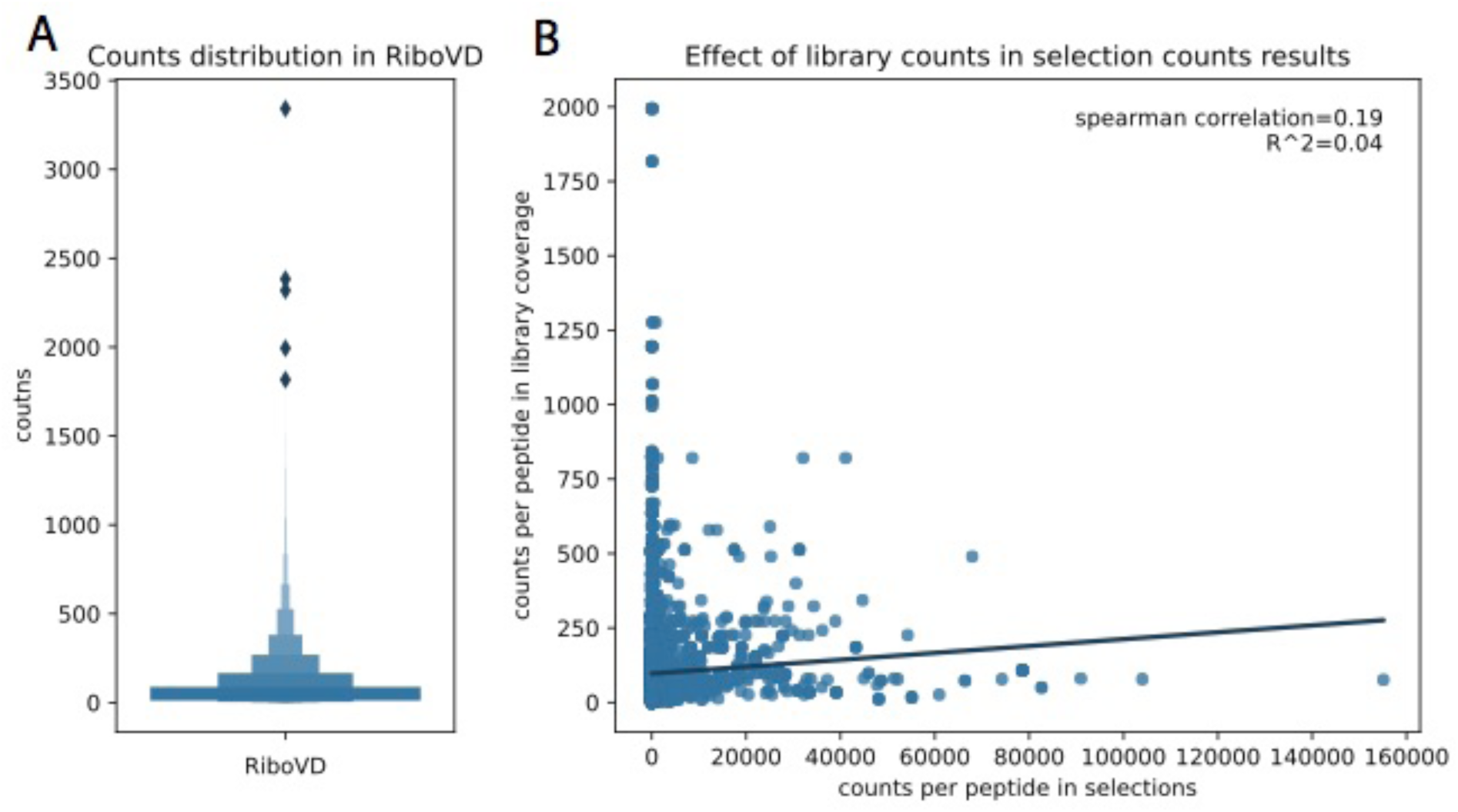
RiboVD counts distribution and effect on selections. **(A)** RiboVD peptides count distribution (sequencing of naive library). **(B)** Correlation analysis between RiboVD peptides counts in coverage vs. medium-/high-confidence peptides counts in selections.

**Figure S2.**
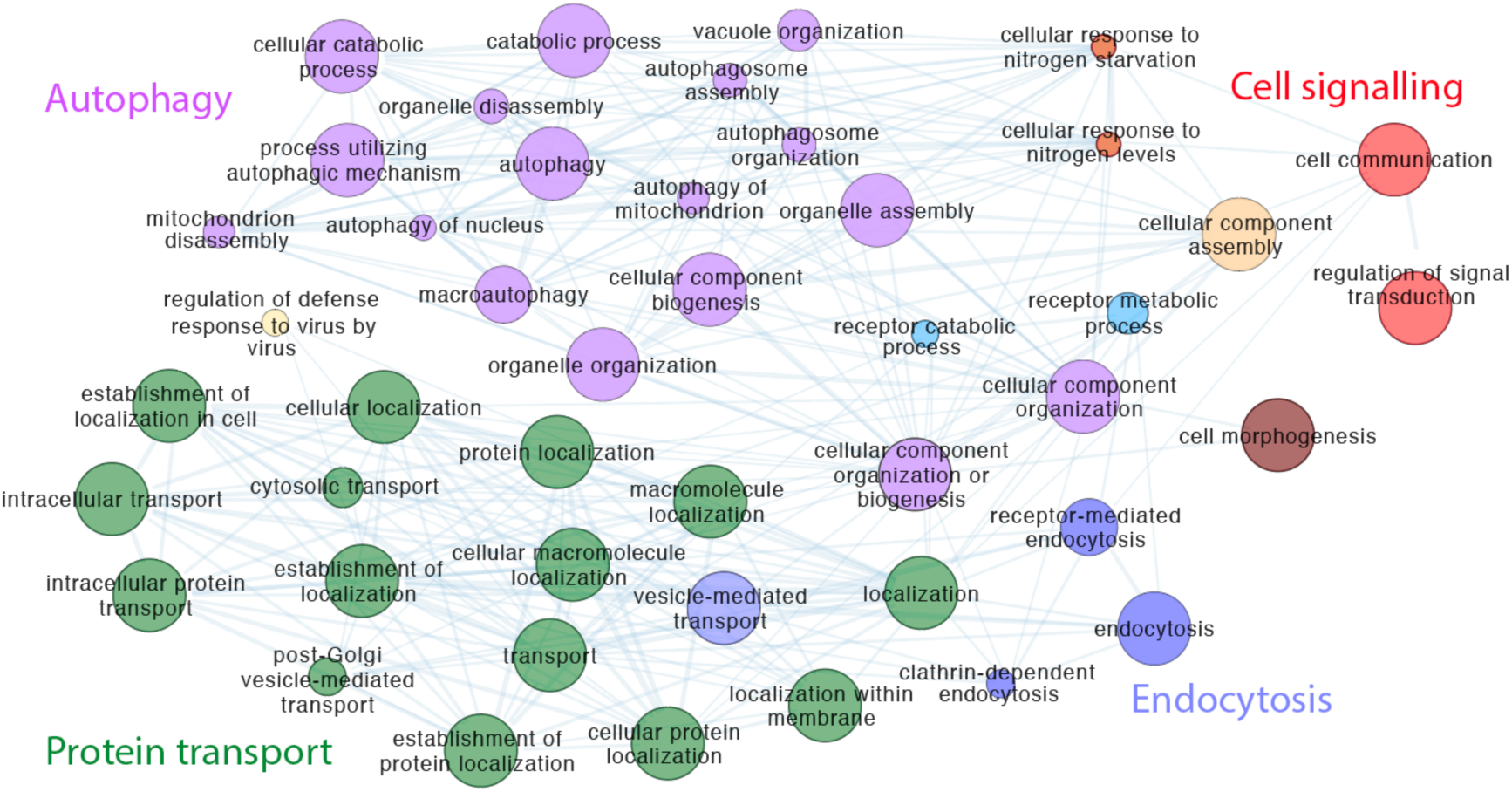
GO Biological Process term enrichment of the expanded host-virus interaction network. Edges represent overlapping genes between the terms, Node size increases with the percentage of genes annotated with the specific term are found in the network, colors correspond to the respective annotated processes on the figure.

**Figure S3.**
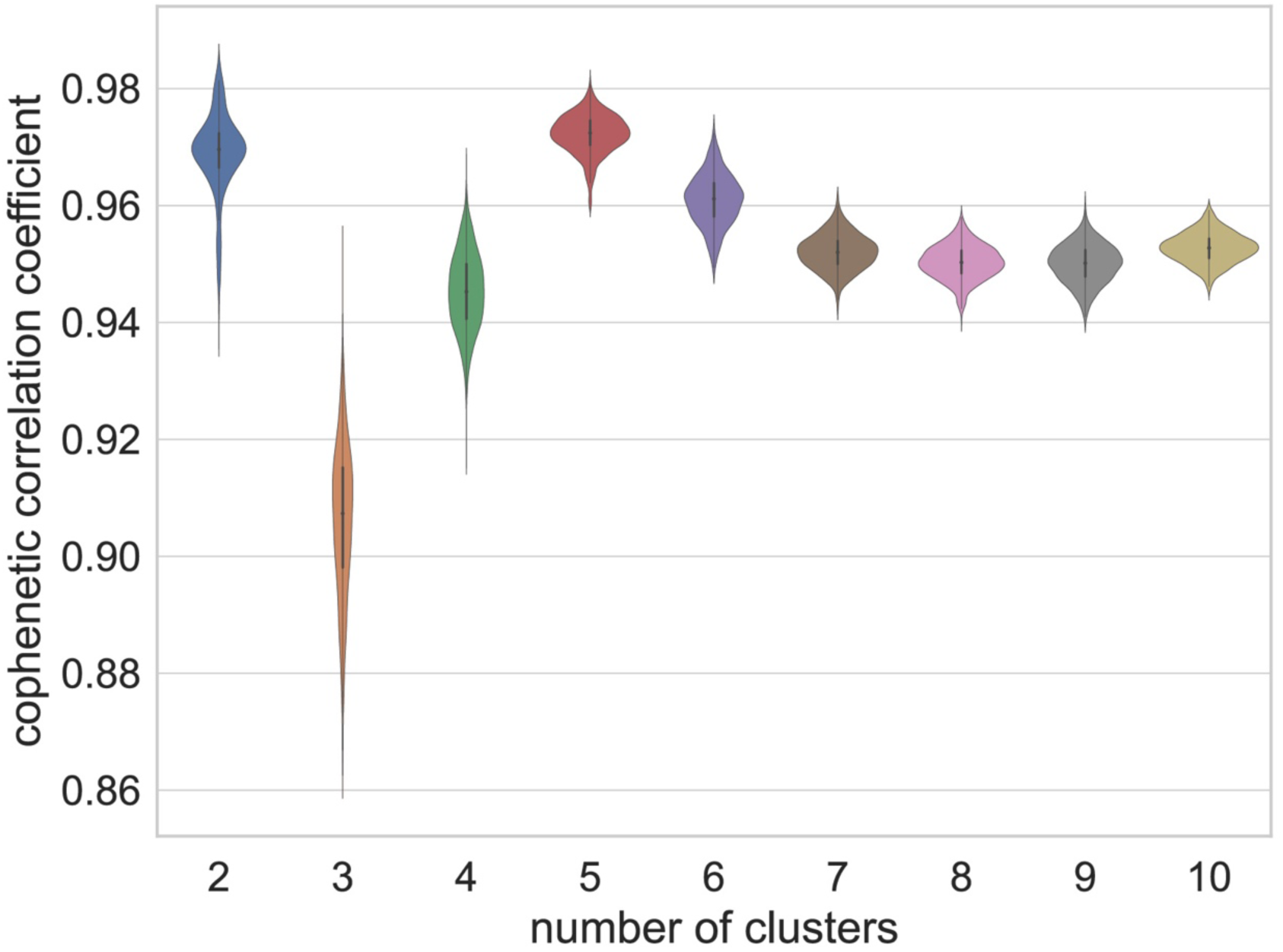
Cophenetic correlation coefficient for different numbers of clusters (latent factors) to determine the numbers of clusters that provides robust cluster membership.

**Figure S4.**
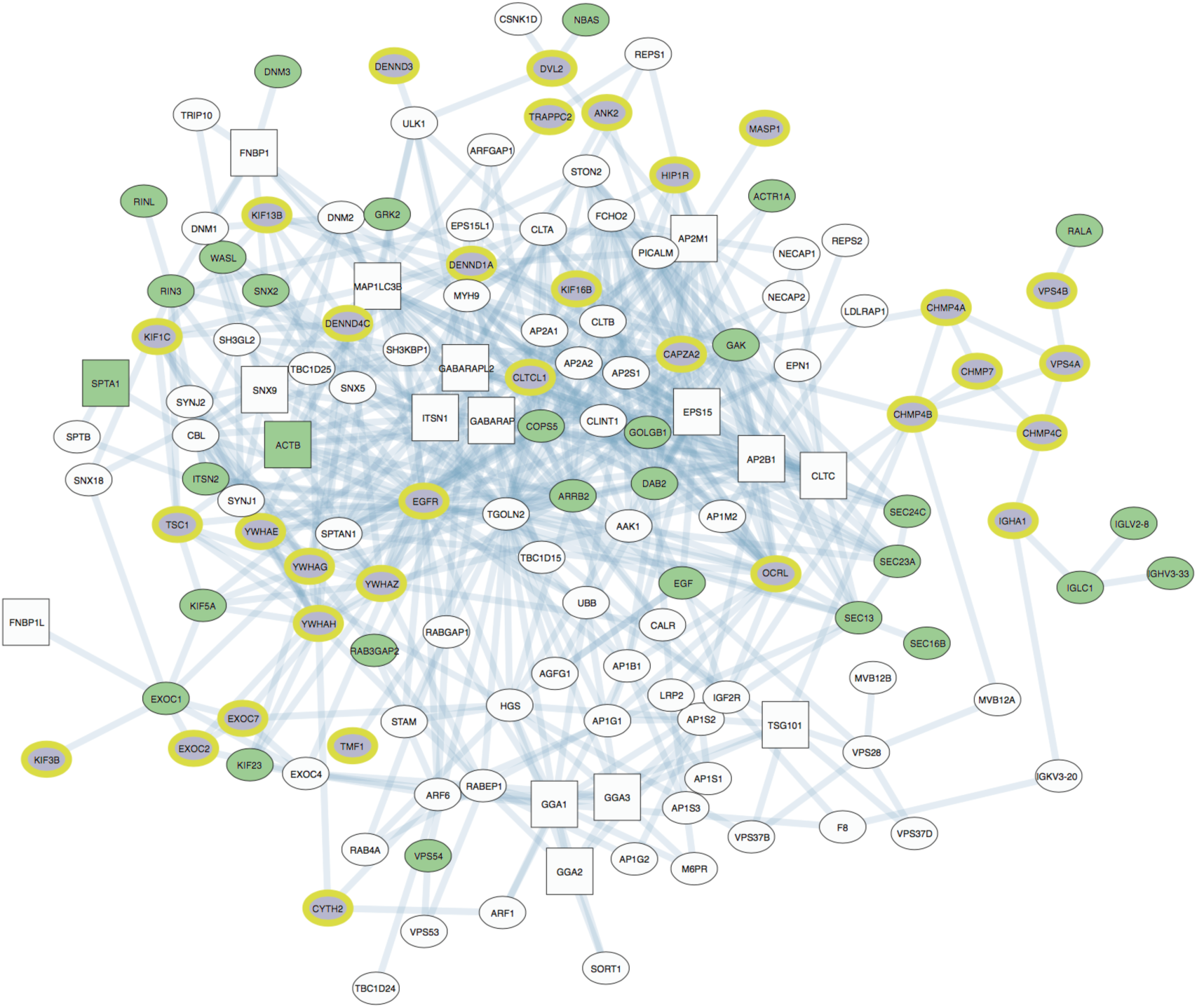
Differences in the vesicle-mediated transport networks targeted by viruses in cluster 4 (green) vs those in cluster 5 (lilac/yellow). White nodes are common to both target networks. Squares indicate proteins used as baits in the experiment.

**Figure S5.**
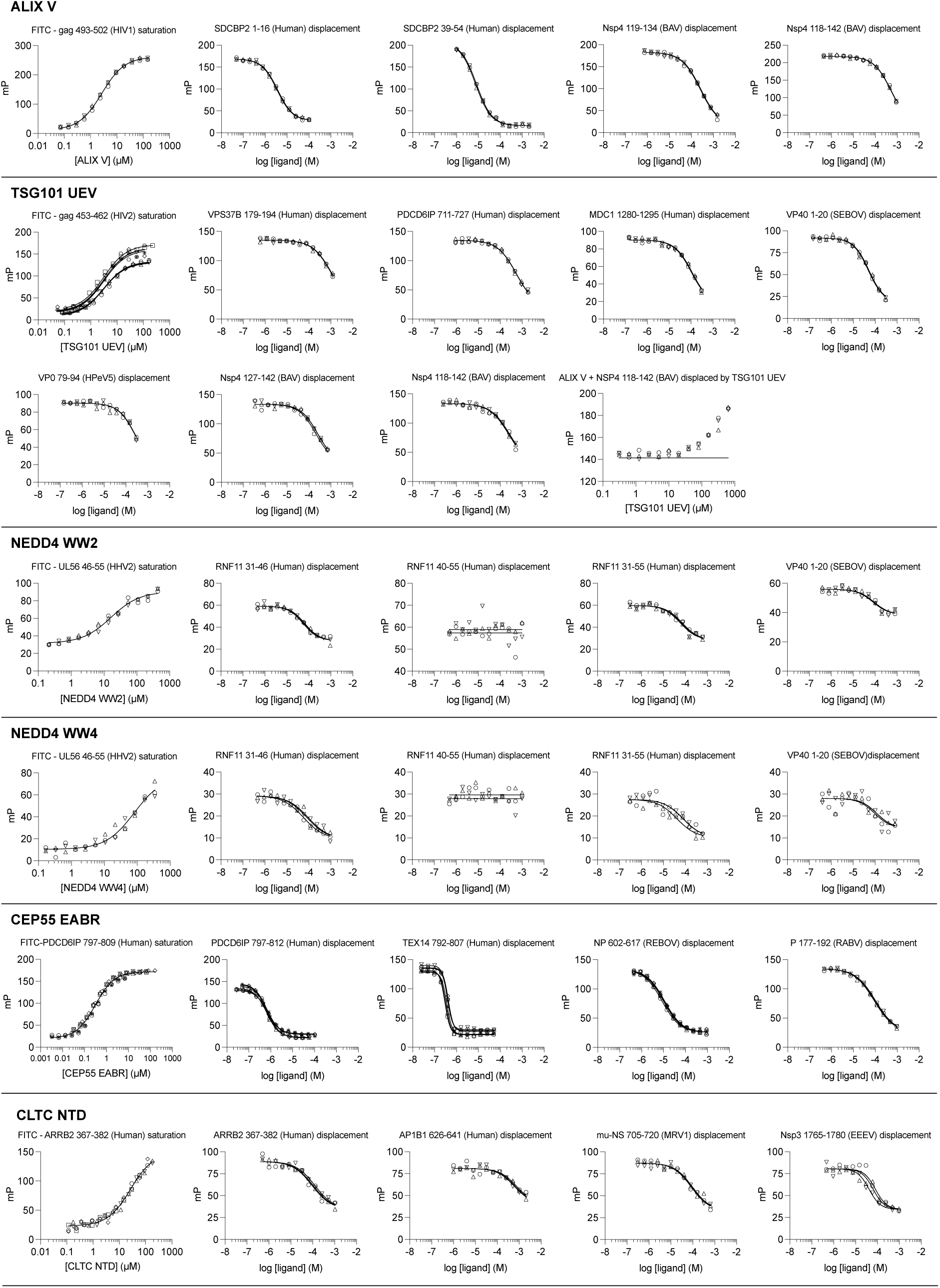
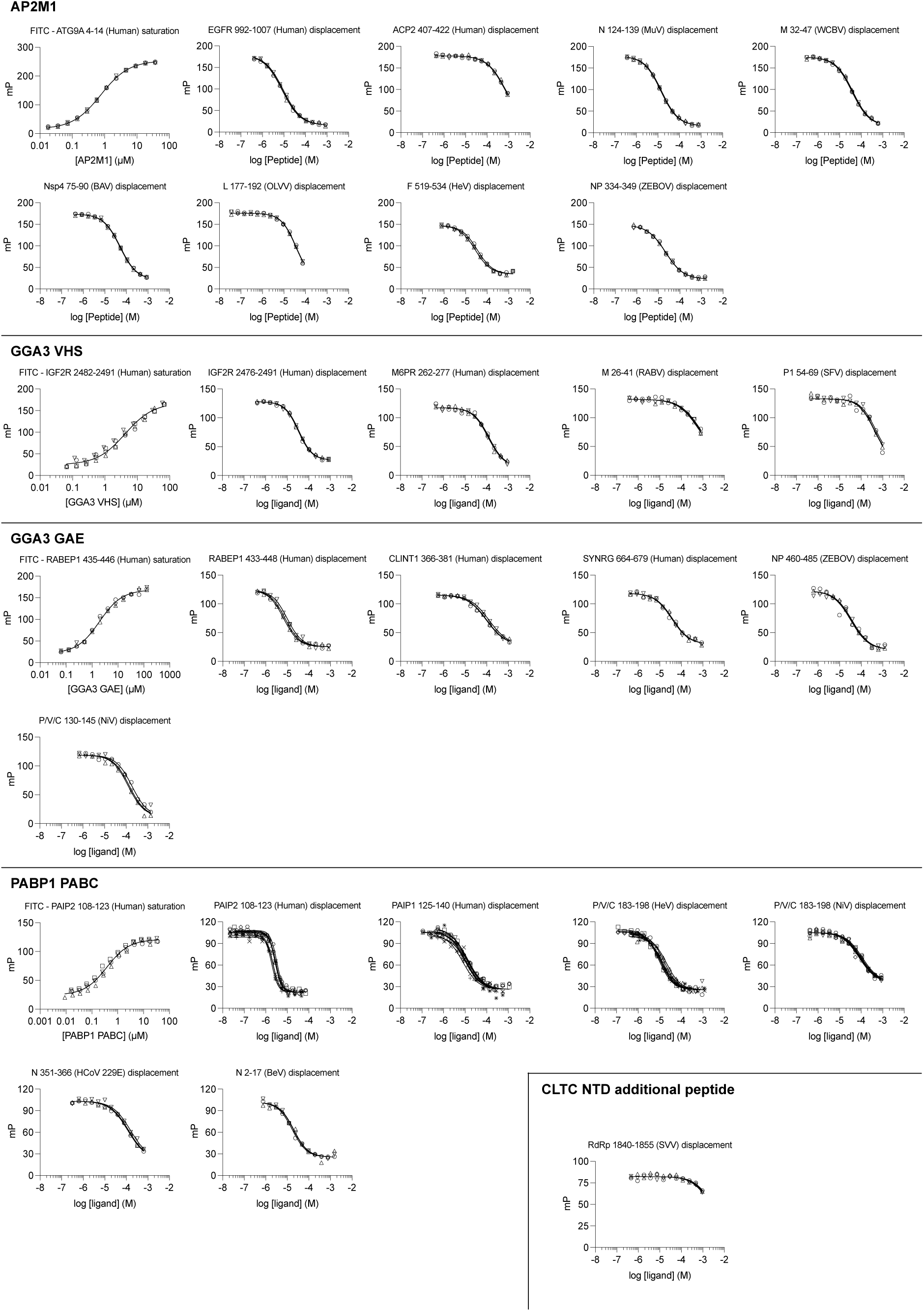
All FP affinity measurements performed in this study. Peptides used, and calculated affinities are presented in Table S7.

**Figue S6.**
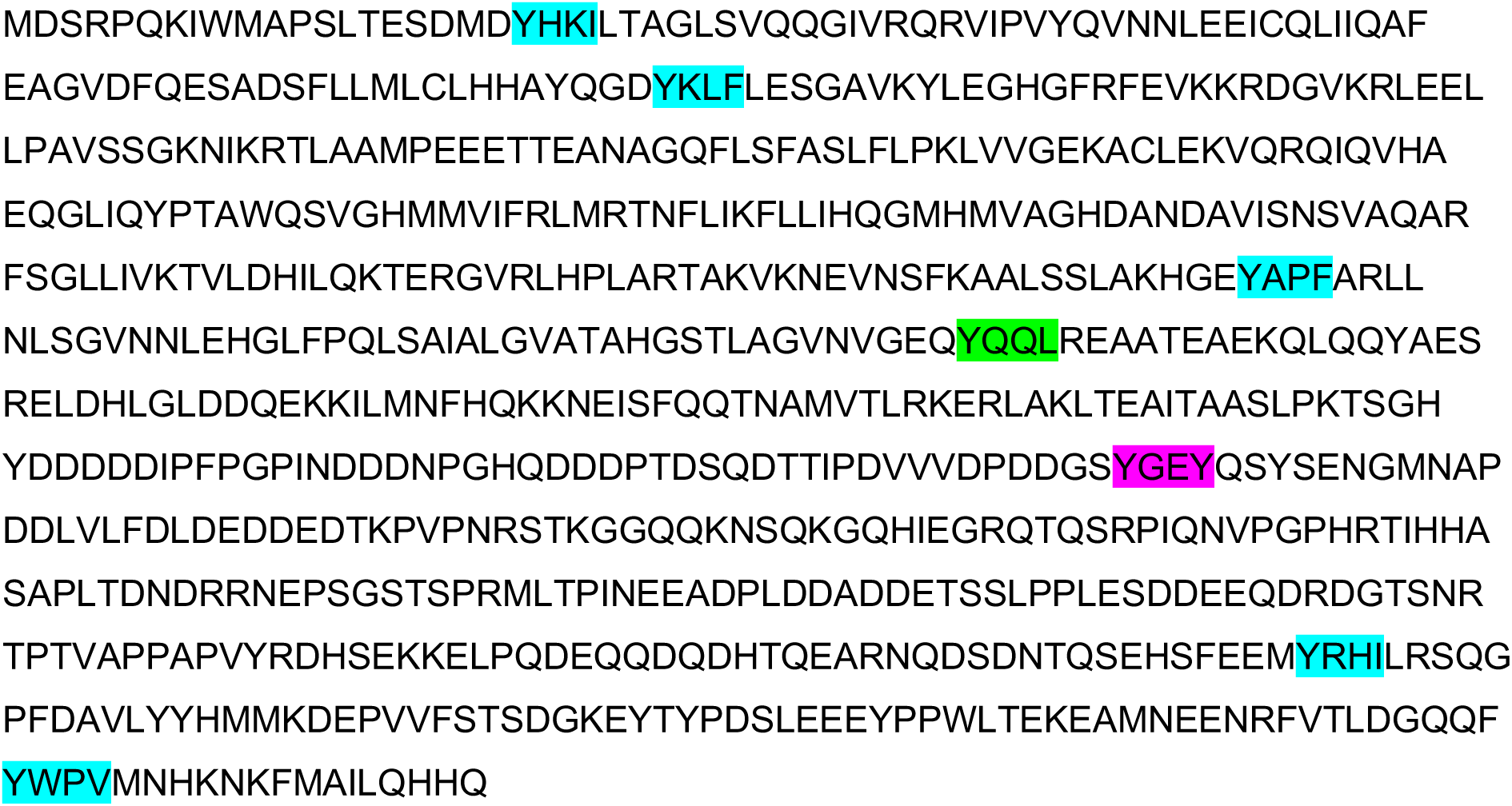
Zaire ebolavirus Nucleoprotein sequence (Uniprot entry P18272). All of the instances matching the AP2M1 recognition motif as reported in eukaryotic linear motif database (ELM) are highlighted in blue. The AP2M1 interaction motif identified in this study is highlighted in green and the GGA3 GAE motif is highlighted in magenta.

**Figure S7.**
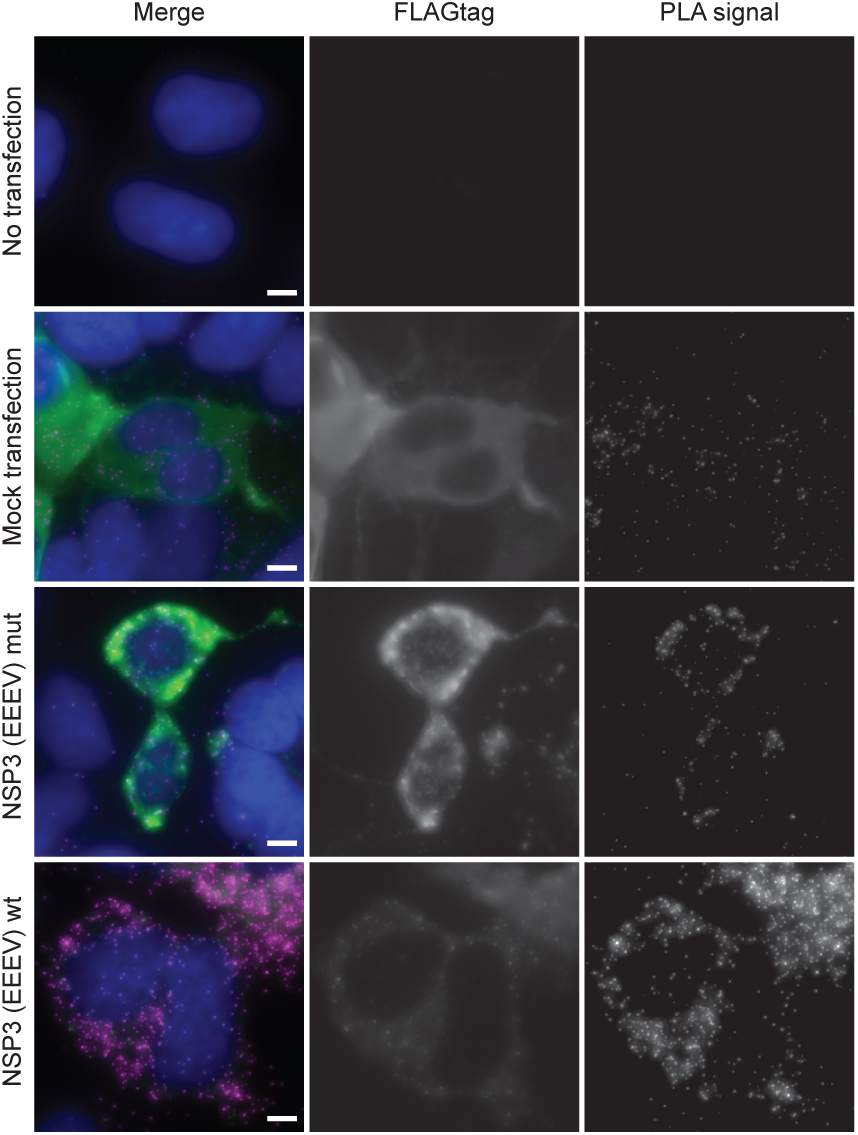
Proximity ligation assay probing the interaction between endogenous clathrin and full length FLAG-tagged NSP3 (EEEV) in HEK293 cells. Fluorescence microscopy images. Nuclei are in blue, FLAG-tag in green and PLA signals visualizing clathrin-NSP3(EEEV) interaction in magenta. Scale bar is 5 μm.

**Figure S8.**
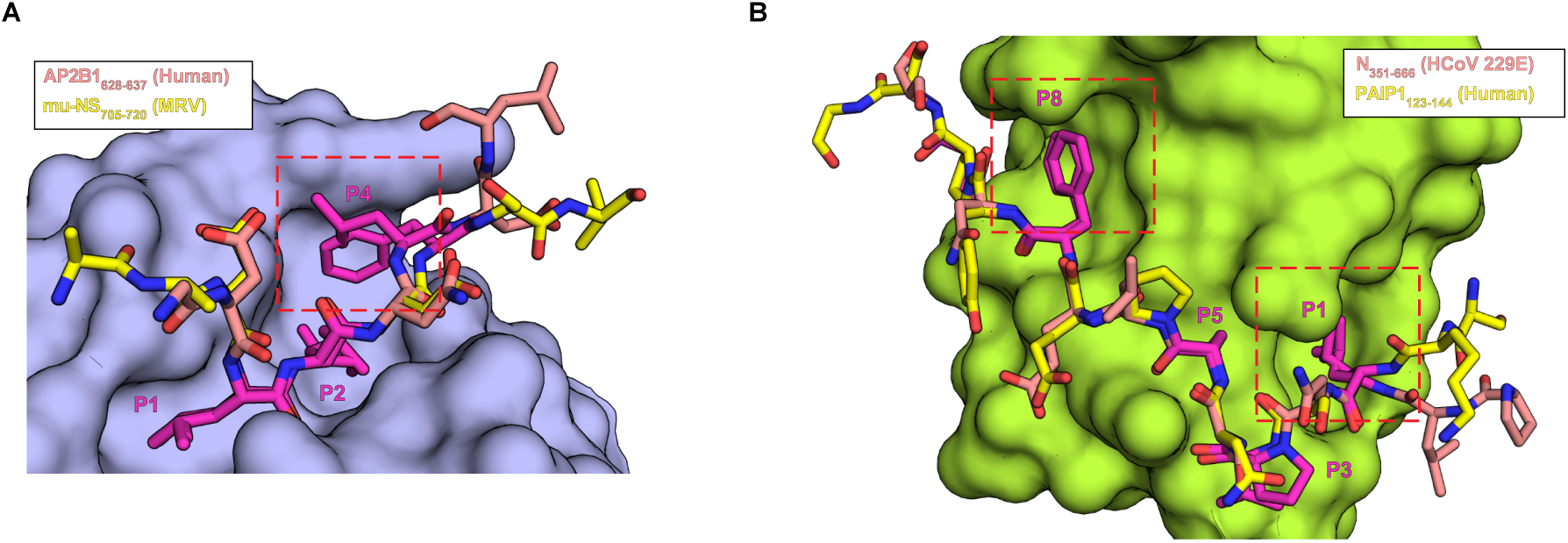
Viral and human ligands bind the CLTC NTD and PABP1 PABC in very similar conformation. **(A)** Crystal structure model of CLTC NTD with superimposed viral ligand from mu-NS (MRV) in yellow and the human ligand from AP2B1 (PDBid: 5M5R) in salmon. The red rectangle highlights the different position of Phe and Leu in position P4. **(B)** Crystal structure model of PABP1 PABC with superimposed viral ligand from N (HCoV 229E) in salmon and the human ligand from PAIP1 (PDBid: 3NTW) in yellow. The red rectangles highlights the two hydrophobic binding pockets at position P1 and P8

**Figure S9.**
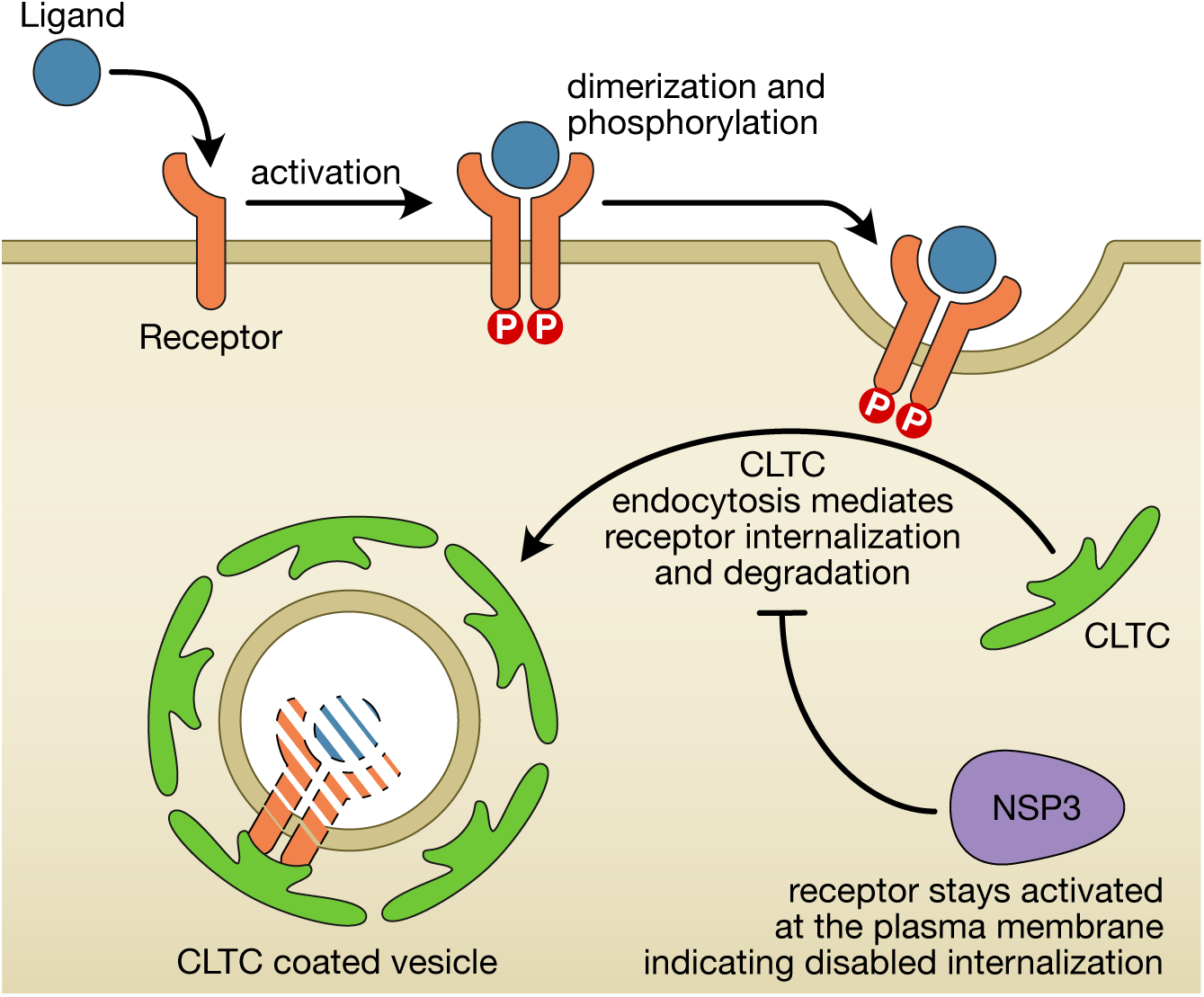
Generalized model of the NSP3 (EEEV) effect on receptor internalization. Upon ligand binding the receptor dimerizes, is activated via autophosphorylation and subsequently internalized from the plasma membrane via clathrin mediated endocytosis. The presence of NSP3 (EEEV) causes the receptor to remain at the plasma membrane in its activated form indicating that clathrin-mediated endocytosis is not functioning properly.

**Figure S10.**
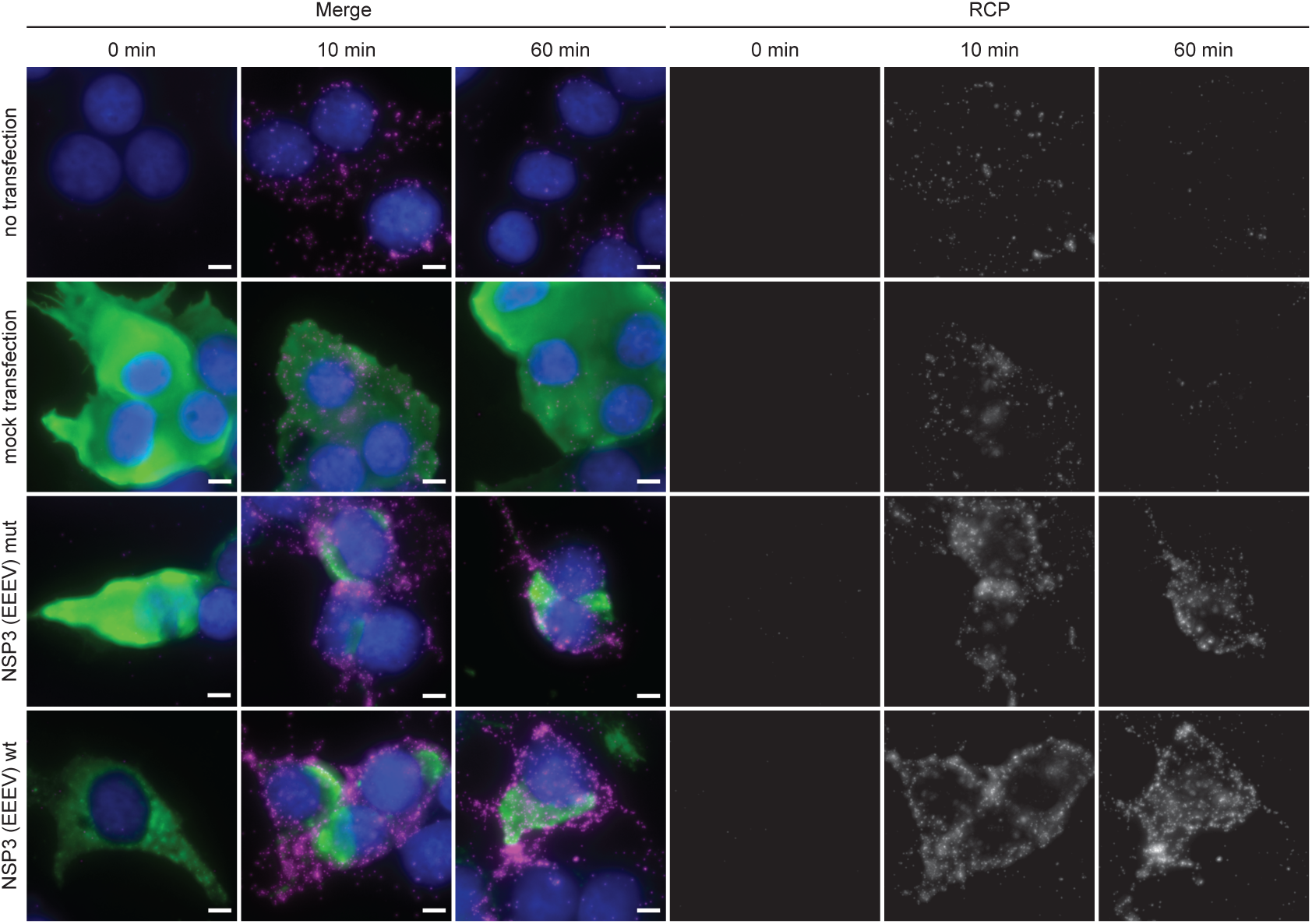
Proximity ligation assay probing the activation of PDGFRβ in HEK293-PDGFRβ-HA cells. Fluorescence microscopy images. Nuclei are in blue, FLAG-tag in green, and PLA signals visualizing phosphorylated PDGFRβ in magenta. Scale bar is 5 μm.

**Figure S11.**
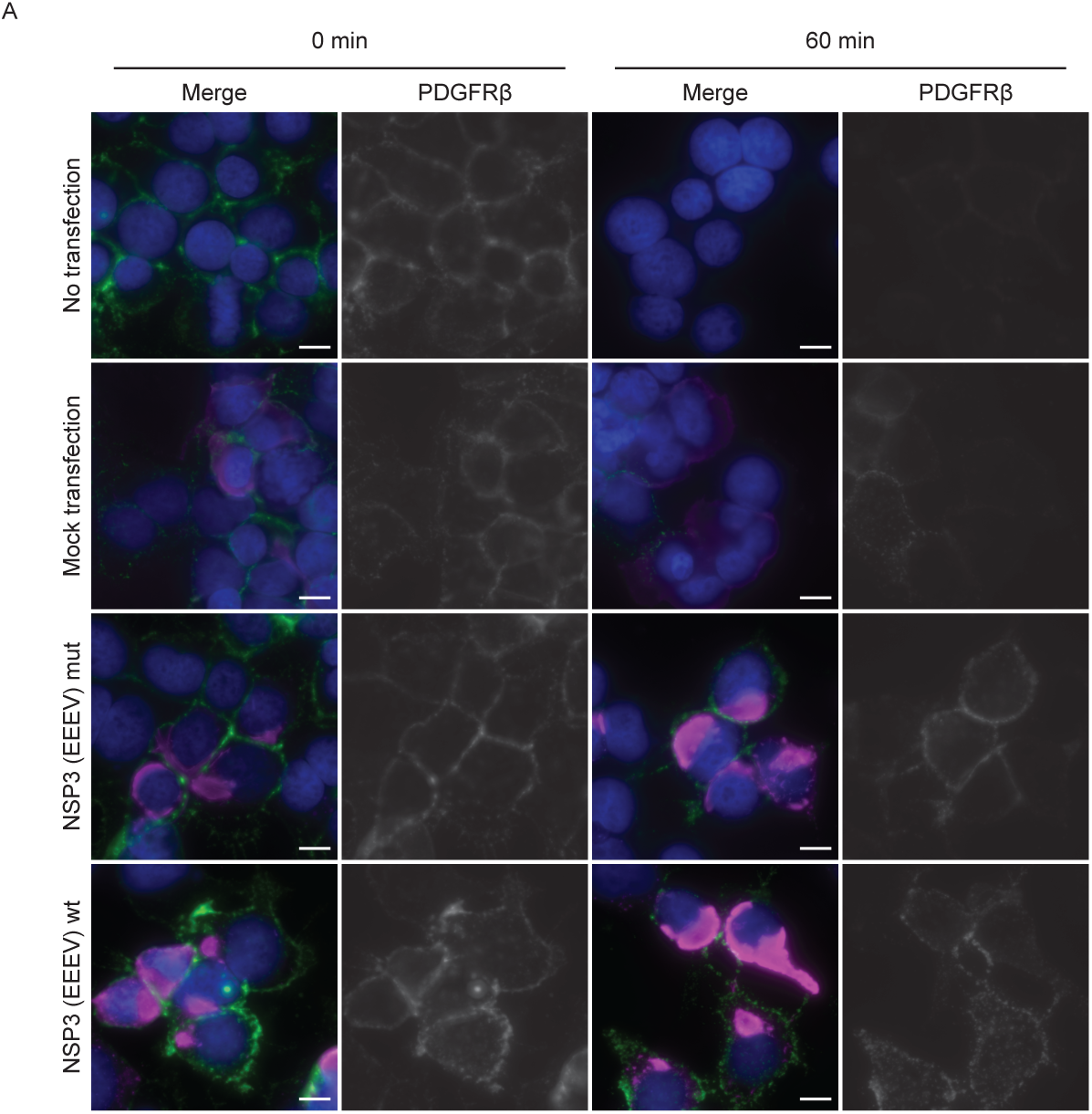
Cell surface fluorescence probing the extracellular part of PDGFRβ. A) Fluorescence microscopy images. Nuclei are in blue, FLAG-tag is in magenta, and PDGFRβ is in green. Scale bar is 5 μm.

**Figure S12.**
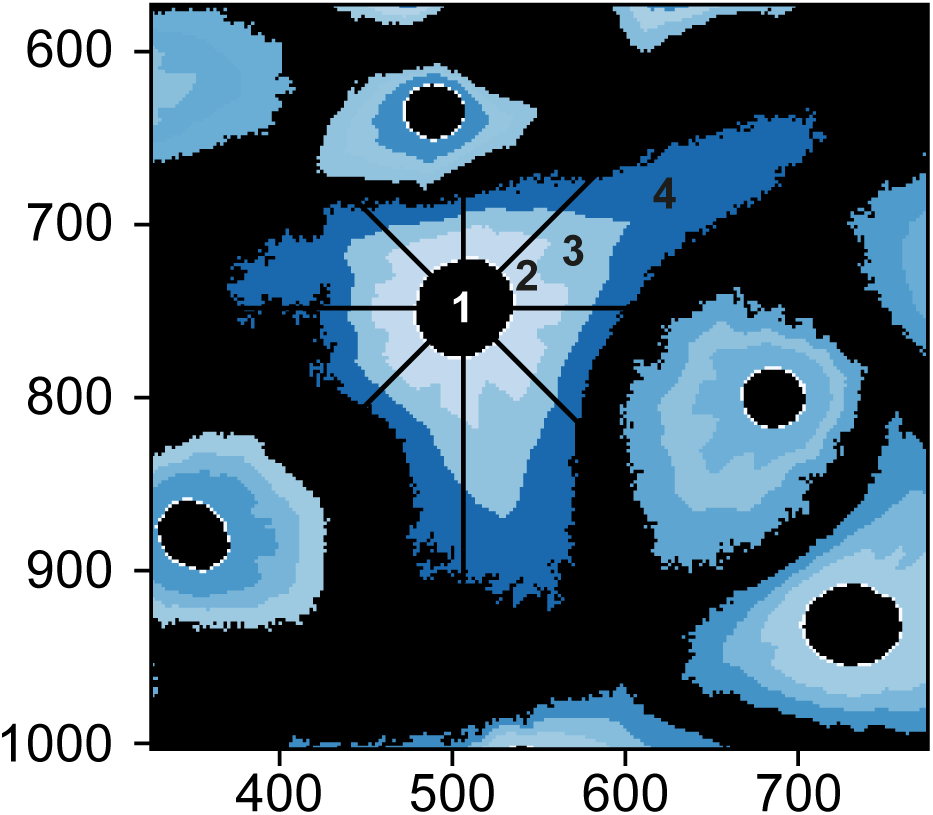
Schematic representation of the measurement of signal distribution and localization in the cell shown in Figure 6J. The distance and radius of the dsRNA replication complexes are detected and grouped in 4 fractions.

**Figure S13.**
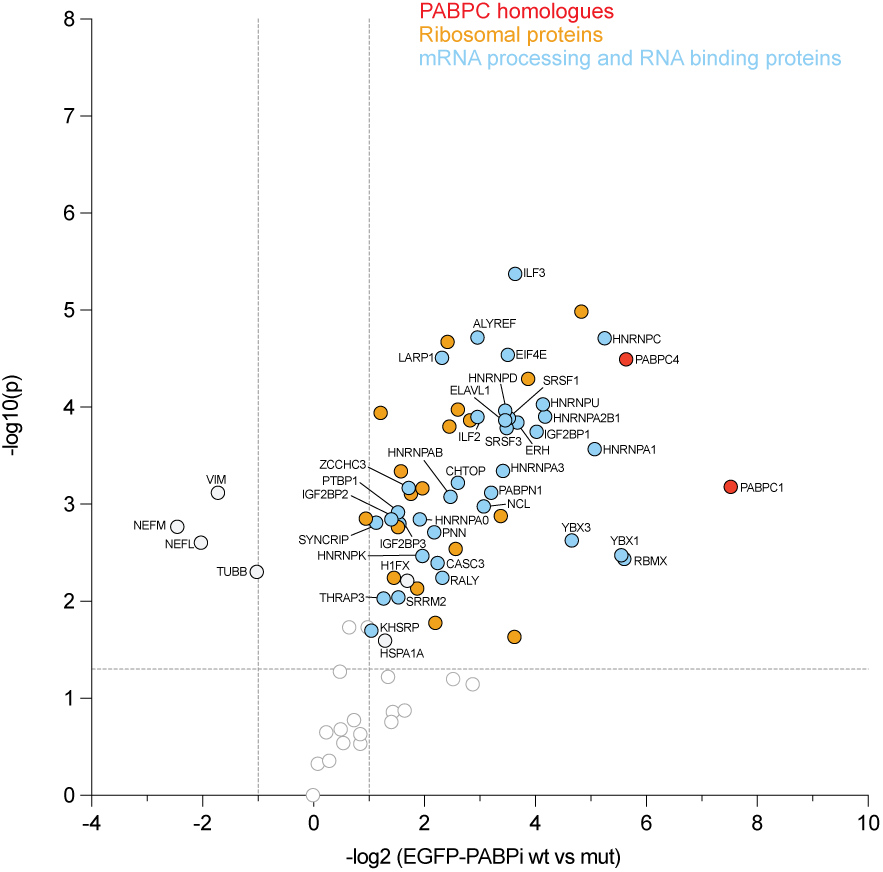
AP-MS results of EGPP-PABPi pulldown from uninfected HEK293 cells.

