## Supplementary material for "Large-scale phage-based screening reveals extensive pan-viral mimicry of host short linear motifs": Table S9 and Table S11

**Table S9**. Overview of crystallographic data.

| | **CLTC-EEV** | **CLTC-mu-NS** | **PAPBC1-HCoV** |
|---|---|---|---|
| **Data Collection** | | | |
| Space group | C2 | C2 | $C222_1$ |
| *a*, *b*, *c* (Å) | 136.9, 129.1, 77.9 | 137.53, 128.9, 78.10 | 63.51, 150.1, 65.5 |
| α, β, γ (°) | 90, 115.3, 90 | 90.00 115.5 90.00 | 90.0 90.0 90.0 |
| Molecules in a. u. | 2 | 2 | 3 |
| Wavelength (Å)[a] | 0.9795 | 0.9795 | 1.003 |
| Resolution (Å) | 70.41-1.96 (1.96-2.11) | 70.49-1.97 (1.97-2.11) | 43.63-1.93 (1.93-2.0) |
| Total reflections | 214797 (10220) | 216228 (10833) | 312878 (31260) |
| Unique reflections | 62270 (3115) | 61571 (3080) | 23731 (2268) |
| Multiplicity | 3.4 (3.3) | 3.5 (3.5) | 13.2 (13.8) |
| Completeness (%) | 90.2 (36.2) | 93.3 (65.2) | 99.1 (98.5) |
| $< I/\sigma(I) >$ | 10.0 (1.6) | 9.1 (1.4) | 12.7 (1.3) |
| Wilson B factor ($Å^2$) | 23.3 | 43.4 | 53.7 |
| $R_{merge}$ [b] | 0.070 (0.673) | 0.078 (0.815) | 0.098 (0.052) |
| $R_{meas}$ | 0.083 (0.804) | 0.093 (0.964) | 0.106 (0.056) |
| $R_{pim}$ | 0.044 (0.434) | 0.049 (0.512) | 0.030 (0.016) |
| $CC_{1/2}$ | 0.998 (0.629) | 0.991 (0.561) | 0.999 (0.999) |
| **Refinement** | | | |
| No. of reflection in work set | 59020 | 58624 | 23713 |
| No. of reflection in free set | 3160 | 2950 | 1124 |
| $R_{work}$ [c] | 0.1800 | 0.1854 | 0.1980 |
| $R_{free}$ [d] | 0.2170 | 0.2189 | 0.2378 |
| No. of non-hydrogen atoms | | | |
| total | 6354 | 6293 | 2268 |
| Protein | 5687 | 5691 | 1858 |
| Solvent | 506 | 470 | 119 |
| Peptide | 161 | 132 | 269 |
| Other ligands | - | - | 19 ($SO_4$) |
| RMS deviations | | | |
| bonds (Å) | 0.009 | 0.013 | 0.007 |
| angles (°) | 1.57 | 1.3 | 0.788 |
| Residues in Ramachandran plot regions (%) | | | |
| favored | 98.49 | 99.0 | 99.3 |
| allowed | 1.51 | 1.0 | 0.4 |
| outliers | 0.00 | 0.00 | 0.4 |
| Average B factor ($Å^2$) | 44.33 | 55.68 | 53.12 |
| Protein | 35.81 | 41.93 | 48.5 |
| Solvent | 40.13 | 46.35 | 56.42 |
| Peptide | 42.58 | 49.8 | 54.46 |
| PDB-ID | 7BN2 | 7BN1 | 7BN3 |

[a] Values in parentheses are for the highest-resolution shell

[b] $R_{merge} = \sum |I - \langle I \rangle| / \sum (I)$ I where is the observed integrated intensity, $\langle I \rangle$ is the average integrated intensity obtained from multiple measurements, and the summation is over all observed reflections.

[c] where

$$R\text{-}factor = \frac{\sum_{(h,k,l)} \| F_{obs}(h,k,l)| - |F_{calc}(h,k,l)\|}{\sum_{(h,k,l)} |F_{obs}(h,k,l)|}$$

$F_{obs}$ and $F_{calc}$ are observed and calculated structure factors respectively.

[d] $R_{free}$ was calculated as for $R_{work}$ but only 5% data left out of refinement procedure has been used in the calculation

### Table S14. Reagent and resource table

| Reagent/resource | Reference or source | Identifier or catalog number |
|---|---|---|
| **Mammalian cells, bacteria, viral strains** | | |
| HEK293 | Sigma | 85120602 |
| HEK293T | TakaraBio | Z2180N (632180) |
| HEK293-PDGFRβ-HA | Kind gift from Frank Böhmer | PMID: 12614164<br>PMID: 10826494 |
| VeroB4 | Kind gift of Gerhard Dobler Bundeswehr Institute of Microbiology, Munich, Germany | |
| VeroE6 | Kind gift of Mattias Forsell Umeå University | |
| SARS CoV-2 | Public Health Agency of Sweden | SARS-CoV-2/01/human2020/SWE accession no/GeneBank no MT093571.1 |
| JEV | Public Health Agency of Sweden | Nakayama strain |
| WNV | Public Health Agency of Sweden | WNV_0304h_ISR00 |
| YFV | Public Health Agency of Sweden | Asibi |
| DENV | Public Health Agency of Sweden | serotype-2; PNG/New Guinea C |
| TBEV | | Torö-2003, PMID: 27982069 |
| LGTV | Kind gift of Gerhard Dobler Bundeswehr Institute of Microbiology, Munich, Germany | TP21 |
| ZIKV | Kind gift of Gerhard Dobler Bundeswehr Institute of Microbiology, Munich, Germany | MR766 |
| RVFV | | Katushka, PMID: 29386590 |

| | | |
|---|---|---|
| **VSV** | **Kind gift of Friedemann Weber, University of Freiburg** | |
| **SINV** | **Kind gift of Olivia Wesula Luande and Magnus Evander** | **Lovanger, KF737350** |
| **CHIKV** | **Kind gift of Magnus Evander** | **CHIKV LR2006OPY1** |
| **E.coli OmniMAX** | **Thermo Fisher Scientific** | **C854003** |
| **E.coli gold BL21 (DE3)** | **Agilent technology** | **230132** |
| **NEB® Stable Competent *E. coli*** | **New England Biolabs** | **C3040I** |
| **Recombinant DNA** | | |
| **pLJM1-EGFP** | **David Sabatini lab** | **Addgene plasmid #19319** |
| **psPAX2** | **Didier Trono lab** | **Addgene plasmid #12260** |
| **pMD2.G** | **Didier Trono lab** | **Addgene plasmid #12259** |
| **Antibodies** | | |
| **Goat-anti-FLAGtag** | **abcam** | **ab1257** |
| **Mouse-anti-FLAG M2** | **Sigma Aldrich** | **F1804** |
| **mouse-anti-clathrin** | **abcam** | **ab2731** |
| **Goat-anti-GST** | **Cytiva** | **274577012** |
| **rabbit-anti-PDGFRβ** | **Cell Signaling Technology** | **#3169** |
| **mouse-anti-PDGFRβ-pY751** | **Cell Signaling Technology** | **#3166** |
| **Donkey anti-Goat IgG (H+L) Highly Cross-Adsorbed Secondary Antibody, Alexa Fluor Plus 647** | **Invitrogen** | **A32849** |
| **Donkey anti-Rabbit IgG (H+L) Highly Cross-Adsorbed Secondary Antibody, Alexa Fluor Plus 555** | **Invitrogen** | **A32794** |
| **goat-anti PDGFRβ** | **RnD Systems** | **AF385** |
| **rabbit-anti-FLAGtag** | **Cell Signaling Technology** | **#14793S** |
| **rabbit anti-SARS-CoV-2 Nucleocapsid** | **Sino Biological Inc** | **40143-R001** |
| **mouse anti-TBEV E 19/1786** | | **pmid 7817895** |
| **mouse anti-Flavivirus Group Antigen** | **ATCC** | **HB-112** |

| | | |
|---|---|---|
| **Antibody, clone D1-4G2-4-15** | | |
| **mouse anti-YFV E** | **ATCC** | **CRL 1689** |
| **mouse anti dsRNA J2** | **Scicons** | **10010500** |
| **rabbit anti-VSV-G** | **Sigma** | **V4888-200UG** |
| **donkey anti-Rabbit IgG (H+L) Highly Cross-Adsorbed Secondary Antibody, Alexa Fluor 555** | **Invitrogen** | **A-31572** |
| **donkey anti-Mouse IgG (H+L) Highly Cross-Adsorbed Secondary Antibody, Alexa Fluor 555** | **Invitrogen** | **A31570** |
| **Goat anti-Rabbit IgG (H+L) Secondary Antibody, HRP** | **Invitrogen** | **31460** |
| **Goat anti-Mouse IgG (H+L) Secondary Antibody, HRP** | **Invitrogen** | **31430** |
| **IRDye® 680RD Donkey anti-Goat IgG secondary Antibody** | **LI-COR** | **925-68074** |
| **IRDye® 800CW Goat anti-Mouse IgG Secondary Antibody** | **LI-COR** | **925-32210** |
| **Oligonucleotides and sequence-based reagents** | | |
| **Chemicals, enzymes and other reagents** | | |
| **Pierce™ Glutathione Agarose** | **Thermo Scientific** | **16100** |
| **Ni sepharose™ excel** | **Cytiva** | **17371201** |
| **$KH_2PO_4$** | **Alfa Aesar** | **7778-77-0** |
| **$K_2HPO_4$** | **VWR** | **7758-11-4** |
| **Peptone** | **Sigma Aldrich** | **91249** |
| **Yeast extract** | **Merck Millipore** | **1.03753.0500** |
| **DTT** | **Fisher bioreagents** | **3483-12-3** |
| **NaCl** | **VWR** | **27810.295** |
| **TCEP** | **Thermo Scientific** | **20491** |
| **Isopropyl-beta-D-thiogalactopyranoside** | **Biosynth Carbosynth** | **367-93-1** |
| **tris(hydroxymethyl)aminomethane** | **Merck Millipore** | **1.08382.1000** |
| **Imidazole** | **Merck Millipore** | **1.04716.1000** |
| **Kanamycin Sulfate** | **VWR** | **25389-94-0** |
| **Ampicillin** | **MEDA/Apoteket** | **011406** |

| | | |
|---|---|---|
| **L-glutathion, reduced** | **Alfa Aesar** | **A18014.06** |
| **Mini-Protean TGX Stain-free gels** | **Bio Rad** | **4568096** |
| **RNase** | **Roche** | **10109134001** |
| **DNase I** | **Roche** | **10104159001** |
| **0.2 µm sterile filter** | **Sarstedt** | **83.1826.001** |
| **PreScission protease** | **Produced in-house** | |
| **Thrombin** | **Cytiva** | **27084601** |
| **M13KO5 helper phage** | **ThermoFisher** | **18311019** |
| **50 bp marker** | **Thermo Scientific** | **10416014** |
| **GelRed** | **Biotium** | **41003-T** |
| **QIAquick PCR Purification Kit** | **Qiagen** | **28104** |
| **Mag-Bind® TotalPure NGS** | **Omega Bio-Tek** | **M1378-00** |
| **DMEM with GlutaMAX Supplement** | **Gibco** | **61965026** |
| **DMEM/F-12** | **Gibco** | **11320033** |
| **FBS** | **Gibco** | **26140087** |
| **MEM Non-Essential Amino Acids Solution** | **Gibco** | **11140035** |
| **Penicillin-Streptomycin** | **Gibco** | **15070063** |
| **Lipofectamine 3000** | **Invitrogen** | **L3000008** |
| **Turbofect Transfection Reagent** | **Thermo Scientific** | **R0531** |
| **Polyethyleneimine, linear** | **Thermo Scientific** | **43896** |
| **DPBS** | **Gibco** | **14190094** |
| **NP-40 Substitute** | **Sigma** | **74385** |
| **cOmplete, EDTA free protease inhibitor** | **Roche** | **05056489001** |
| **DC assay kit** | **Bio-Rad** | **5000114/5000113/5000115** |
| **GFP-Trap Dynabeads** | **Chromotek** | **gtd-20** |
| **Lysozyme** | **ITW reagents** | **A4972** |
| **Phusion High-Fidelity polymerase** | **Thermo Fisher Scientific** | **F-530S** |
| **Trypsin** | **Promega** | **V5111sn** |
| **C18 membrane** | **3M Empore** | **2215** |
| **Opti-MEM** | **Gibco** | **11058021** |
| **Intercept (TBS) Blocking Buffer** | **Li-cor** | **927-60001** |
| **TrueBlue peroxidase substrate** | **KPL** | **KPLI50-78-02** |
| **Duolink PLA probe anti-Mouse PLUS** | **Olink** | **82021** |

| | | |
|---|---|---|
| **Duolink PLA probe anti-Goat MINUS** | **Olink** | **82006** |
| **Duolink PLA probe anti-Rabbit MINUS** | **Olink** | **82005** |
| **Duolink Ligation solution** | **Olink** | **82009** |
| **Duolink Amplification Red solution** | **Olink** | **82018** |
| **T4 DNA ligase** | **Thermo Scientific** | **EL0016** |
| **Phi29 DNA polymerase** | **Thermo Scientific** | **4002** |
| **Recombinant Human PDGF-BB** | **Peprotech** | **100-14B** |
| **Plates and flasks** | | |
| **175 cm2 CellBIND™ Surface Cell Culture Flasks** | **Corning Life Sciences** | **3292** |
| **Nunc MaxiSorp plates** | **Thermo Fisher Scientifc** | **44-2404-21** |
| **Black, non-binding surface, flat bottom 96-well plates** | **Corning Life Sciences** | **3993** |
| **Tissue culture Dish 100mm** | **Sarstedt** | **83.3902** |
| **Nunc™ Lab-Tek™ II CC2™ Chamber Slide System** | **Sigma** | **S6815** |
| **MRC 2 Well Crystallization Plate in UVXPO** | **Hampton Research** | **HR3-106** |
| **Software** | | |
| **MaxQuant** | **2.0.1.0** | **https://www.maxquant.org/ https://doi.org/10.1038/nbt.1511** |
| **Perseus** | **2.0.3.0** | **https://www.maxquant.org/perseus/ https://doi.org/10.1038/nmeth.3901** |
| **Cellprofiler** | **3.0.0** | **https://cellprofiler.org/previous-releases** |
| **Leica Application Suit X software** | | **Leica** |
| **Cytoscape** | **v.3.9.1** | **https://cytoscape.org/download.html** |
| **GraphPad Prism** | **v9.3.1** | **https://www.graphpad.com/scientific-software/prism/** |
| **Pymol** | **v2.3.5** | **https://pymol.org/2/** |
| **Instruments and other** | | |
| **iD5 plate reader** | **Molecular Devices** | |

| | | |
|---|---|---|
| iD5 plate reader | Molecular Devices | |
| Concentrator Plus | Eppendorf | |
| Illumina MiSeq v3 | Illumina | MS-102-3001 |
| Syringe filter, Filtropur S | Sarstedt | 83.1826.001 |
| Easy-nLC 1000 | Thermo Scientific | |
| Zen 2 (Blue edition) | Zeiss | |
| Acclaim PepMap 100 pre-column | Thermo Scientific | 164535 |
| PepMap RSLC C18 analytical column | Thermo Scientific | 164534 |
| Q Exactive Plus | Thermo Scientific | |
| TROPHOS Plate RUNNER HD® | Dioscure, Marseille, France | |
| Plate washer | Molecular devices | |
| Odyssey CLx Imaging system | LI-COR | 9140 |
| MALDI TOF/MS Ultraflex III | Bruker | |
| Diamond Light Source | | Didcot, UK |
| Zeiss Imager Z2 | Zeiss | |
